# A Parsimonious Description of Global Functional Brain Organization in Three Spatiotemporal Patterns

**DOI:** 10.1101/2021.06.20.448984

**Authors:** Taylor Bolt, Jason S. Nomi, Danilo Bzdok, Jorge A. Salas, Catie Chang, B.T. Thomas Yeo, Lucina Q. Uddin, Shella D. Keilholz

**Affiliations:** Emory University/Georgia Institute of Technology, Atlanta, GA, USA; Department of Psychiatry and Biobehavioral Sciences, University of California Los Angeles, Los Angeles, CA, USA; The Neuro - Montreal Neurological Institute (MNI), McGill University & Mila - Quebec Artificial Intelligence Institute, Montreal, QC, Canada; Vanderbilt University, Nashville, TN, USA; Department of Electrical & Computer Engineering, Centre for Translational MR Research, Centre for Sleep & Cognition, N.1 Institute for Health and Institute for Digital Medicine, National University of Singapore, Singapore

**Keywords:** propagation, dynamics, time-lag, global signal, task-positive, task-negative, gradient, resting-state fMRI, taxonomy

## Abstract

Resting-state functional MRI has yielded seemingly disparate insights into large-scale organization of the human brain. The brain’s large-scale organization can be divided into two broad categories - zero-lag representations of functional connectivity structure and time-lag representations of traveling wave or propagation structure. Here we sought to unify observed phenomena across these two categories in the form of three low-frequency spatiotemporal patterns composed of a mixture of standing and traveling wave dynamics. We showed that a range of empirical phenomena, including functional connectivity gradients, the task-positive/task-negative anti-correlation pattern, the global signal, time-lag propagation patterns, the quasiperiodic pattern, and the functional connectome network structure are manifestations of these three spatiotemporal patterns. These patterns account for much of the global spatial structure that underlies functional connectivity analyses, and unifies phenomena in resting-state functional MRI previously thought distinct.

## Introduction

Since the discovery of spontaneous low-frequency blood-oxygenation-level dependent (BOLD) fluctuations in the 1990s^1^, increasingly complex analytic techniques have been applied to understand the spatial and temporal structure of these brain signals. A notable feature of these signals is their organization into global patterns that span across functional systems for cognition, perception and action ^2–4^.

We distinguish between two characterizations of this global structure: zero-lag synchrony and time-lag synchrony between brain regions. Zero-lag synchrony is defined as instantaneous statistical dependence between two time courses, or the correlation between two BOLD signals with no time-lag. The zero-lag analysis approach has identified several global patterns spanning across functional networks that have generated sustained research interest: the global signal ^5^ , the task-positive/task-negative pattern ^6^ , and the principal FC gradient ^2^.

Time-lag synchrony is defined as the statistical dependence between two time courses, where one time-course is delayed in time. Two prominent global patterns with coherent time-lag structure have emerged. Short spontaneous global propagation BOLD fluctuations that extend across cortical and subcortical brain regions (∼0 - 2s) are known as lag projections^7^ . This propagation pattern varies according to experimental manipulations of task demands and sensory inputs, suggesting that at least some of this structure is uncoupled from hemodynamic delays ^7^. A pseudo-periodic spatiotemporal pattern at a longer time-scale (∼20s) known as the ‘quasi-periodic pattern’ (QPP) involves an alteration in BOLD amplitudes between the task-positive (TP) and default mode networks (DMN). The shift in BOLD amplitudes between TP and DMN regions is marked by a large-scale propagation of BOLD activity between the two networks.

There may be an underlying unity to these representations that has heretofore remained overlooked. We hypothesized that the vast majority of widely-used zero-lag and time-lag representations of intrinsic functional brain organization capture different aspects of a small number of spatiotemporal patterns that exhibit *both* zero-lag and time-lag structure. Our specific hypotheses were 1) global patterns of zero-lag and time-lag synchrony are describing different facets of the same underlying spatiotemporal patterns, and 2) a small set of spatiotemporal patterns can explain a large number of previous findings in the literature describing spontaneous BOLD signal fluctuations.

Three lines of evidence support the first hypothesis. First, time-lag representations have a spatial distribution that precisely maps to the spatial weights of the principal FC gradient ^8–10^. Second, the cortical global signal spatial topography is not entirely constituted by zero-lag spatial structure, but has significant time-lag structure ^10,11^. Third, removal of time-lag synchronous patterns, such as the QPP, from spontaneous BOLD fluctuations substantially alters patterns of zero-lag synchrony in FC network representations ^12^. These findings suggest that there may be a common pattern of global BOLD activity that unifies these zero-lag and time-lag representations.

To develop intuitions on the proposed relationship between zero-lag and time-lag synchrony patterns, we utilize the concepts of ‘standing’ and ‘traveling’ waves ^13,14^. Standing waves refer to stationary oscillations exhibiting no time-lagged statistical dependencies across space (of the kind captured by FC analyses). Traveling waves refer to oscillations in a spatial field with non-zero time-lag statistical dependence across space (of the kind captured by the QPP and lag projection algorithm). We suggest that global BOLD spatiotemporal patterns consist of a mixture of standing and traveling wave spatial structure. Zero-lag analyses capture the standing wave structure of these patterns, while time-lag analyses capture the traveling wave structure of these patterns. To capture these patterns in a single latent representation, we utilize a complex-valued extension of a popular dimension reduction technique: complex principal component analysis (CPCA).

In support of the second hypothesis, we begin with the observation that the resting-state fMRI literature reveals very similar patterns of global BOLD activity across analytic approaches, including FC gradients ^2^ , CAPs^15^ , independent component analysis (ICA) ^16^ , and other latent brain state methods ^17^ , as well as seed-based correlation analyses ^6^. We propose that these similar patterns across analysis methods are descriptions of the same underlying spatiotemporal patterns proposed in our first hypothesis. To test the second hypothesis, we compared a systematic survey of zero- and time-lag analyses to a set of spatiotemporal patterns derived from CPCA.

Our analyses revealed that three spatiotemporal patterns constitute the dominant large-scale spatial structure in spontaneous low-frequency BOLD fluctuations. With these three patterns, we can unify a range of previous findings in resting-state fMRI, including lag projections ^7^, the QPP^18^, the topography of the global signal ^19^, the task-positive/task-negative pattern^6^, the principal FC gradient^2^, and FC network structure. We demonstrate that all of these previous observations are manifestations of three spatiotemporal patterns captured within a unifying framework capable of modeling standing and traveling oscillatory BOLD phenomena. This novel framework allows for a parsimonious description of global functional brain organization that can inspire new hypotheses about the mechanisms underlying coordination of spontaneous activity across the brain.

## Results

### 1. Standing and Traveling Wave Simulation

We posit that the relative mixture of standing and traveling waves in cortical BOLD signals explains the spatial similarity between outputs of zero-lag and time-lag analyses. To test this claim, we first conducted a simulation study of varying degrees of traveling and standing spatiotemporal wave patterns.

Standing and traveling wave simulations (see **Supplementary Modeling Note 1**) consisted of a back-and-forth sinusoidal oscillation of Gaussian curves on a two-dimensional grid (**Figure 1**). This approach allowed us to systematically vary the degree of traveling wave behavior in each oscillation by adjusting the distance between Gaussian curve peak locations, from a distance of zero (‘pure standing’ motion), to a large distance (‘pure traveling’ motion). Zero-lag dimension-reduction techniques were applied to the time series of this simulated oscillation.

**Figure 1.**
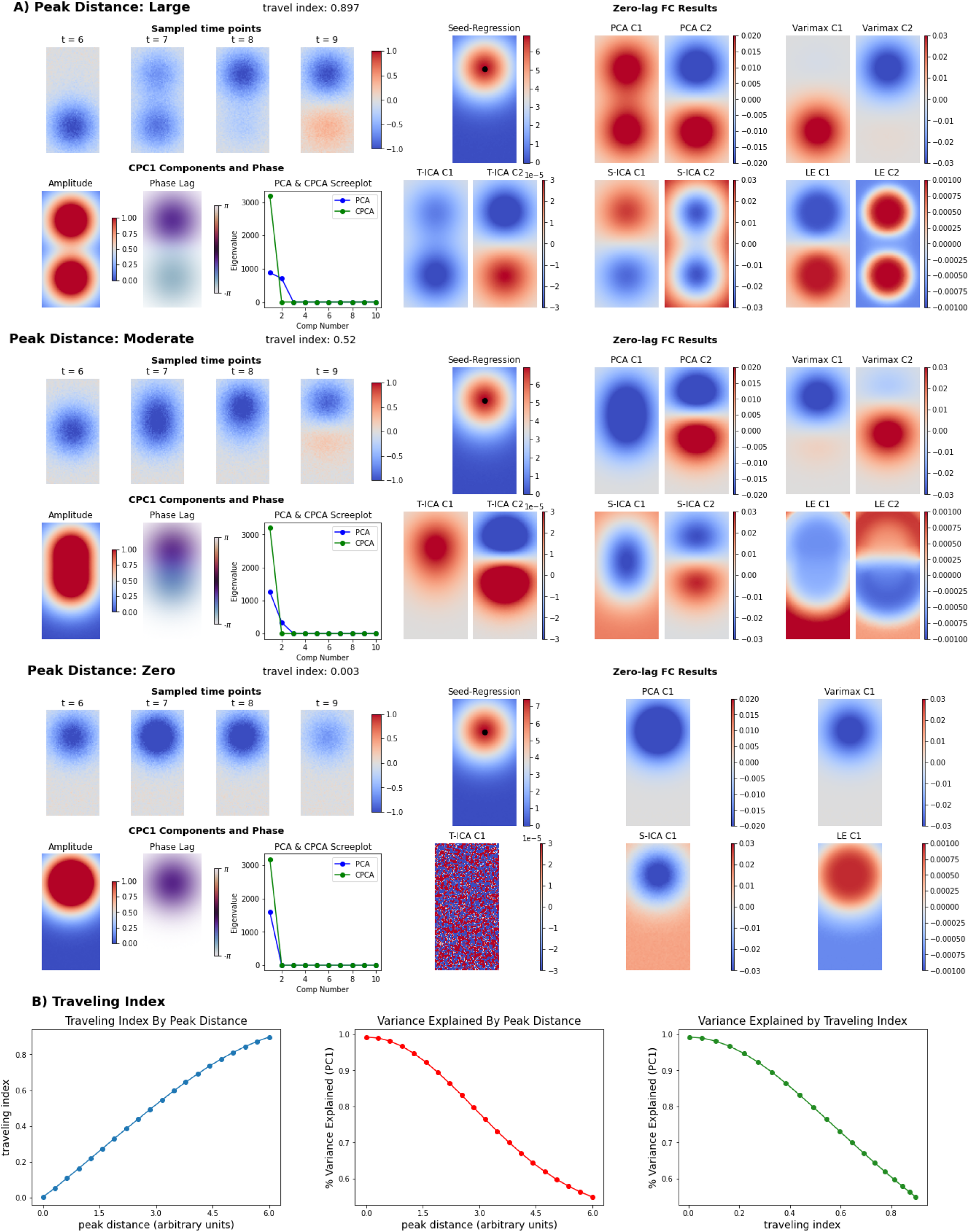
Simulation to Analyze Standing and Traveling Wave Oscillations. **A)** Visualization of simulation and analysis results of traveling two-dimensional Gaussian curves at three peak distances (large, moderate and zero distance). For each simulation scenario, the bottom Gaussian curve moves upward (bottom to top) towards the top Gaussian curve. In the top left panel of each simulation are four arbitrarily sampled time points. Note, the travel distance between the two Gaussian curves grows smaller at smaller peak distances (moving from the top to bottom panel). The amplitude and phase-lag maps of the first complex principal component from CPCA are shown in the bottom left panel of each simulation. For all simulations, the amplitude and phase maps of the first complex component accurately describe the spatial distribution and phase-lag between the two Gaussian curves. In addition, the scree plot of the zero-lag (blue) and complex-valued (green) correlation matrix is displayed. Results of various zero-lag FC analyses are displayed in the right panel. For the dimension-reduction techniques, two components were estimated at non-zero peak distances, motivated by the non-zero second eigenvalue of the correlation matrix. For zero-peak distance, only a single component was estimated. At non-zero peak distances, zero-lag dimension-reduction techniques tend to split the traveling wave oscillation into either 1) separate Gaussian curves (varimax), or 2) into separate phases of the oscillation. **B**) From left to right, plots of the traveling index by peak distance, the variance explained by the first eigenvalue (zero-lag correlation matrix) by peak distance, and the variance explained by the first eigenvalue by traveling index. PCA = principal component analysis; CPCA = complex PCA; S-ICA **=** spatial ICA; T-ICA = temporal ICA; LE = Laplacian eigenmaps

Zero-lag dimension-reduction analyses applied to pure standing waves effectively captured the oscillation in one latent factor (**Figure 1A**; zero peak distance). Using Catell’s scree plot test ^20–22^, we identified the point where the successive extraction of latent components exhibits a flattening in explained variance as the optimal number of components to extract. For pure standing motion, only the first eigenvalue was non-zero, confirming a single latent factor. Seed-based correlation analysis from a seed at the center of one Gaussian curve returned the same spatial pattern.

At non-zero distance between Gaussian curve peak locations, zero-lag analyses separated the non-synchronous motion into two components. The second eigenvalue of the correlation matrix increased with larger distance, indicating the presence of two latent components. The spatial patterns of the two components were largely consistent across methods. However, methods favoring spatial weight sparsity separated the traveling motion into isolated Gaussian curves, while non-sparse methods extracted separate phases or ‘snapshots’ of the overall oscillation.

Dimension-reduction techniques were extended into the complex-valued domain by augmenting the real time-courses of the grid into a complex signal via the Hilbert transform. To demonstrate the utility of complex-valued dimension-reduction methods in extracting traveling wave oscillations, we applied complex-valued PCA (CPCA) to the same empirical data simulations of traveling motion (**Supplementary Modeling Note 1).** Analogous to our diagnostics of the zero-lag correlation matrix, we constructed a scree plot consisting of the ordered eigenvalues from the complex-valued correlation matrix constructed from the complex grid time courses.

For pure traveling motion, the scree plot from the complex-valued correlation matrix revealed that only the first eigenvalue was non-zero, indicating the existence of a single latent factor (**Figure 1A**; large peak distance). The grid amplitudes of the first complex principal component reflected the spatial distribution of the two Gaussian curves, indicating that the coherent fluctuations of the two Gaussian curves were captured by a single latent component. Importantly, the phase-lag values of the first component precisely reflected the back-and-forth motion of the Gaussian curves.

In simulations with non-zero distance between peak locations, CPCA and zero-lag techniques converged on a similar solution. Further, at smaller distances, the scree plots of the zero-lag and complex-valued correlation matrix both yielded a one-factor solution.

By construction, this simulation had access to the ‘ground truth’ mixture of traveling and standing wave components. No such ground truth is available in BOLD signals recorded from the brain. Thus, we invoked a ‘traveling index’ ^14^ that measures the presence of traveling waves in CPCA components, varying from 0 (pure standing waves) to 1 (pure traveling waves). Such a metric provides a quantitative estimate of the mixture of traveling and standing waves in the absence of ground truth simulated data.

From the observation that zero-lag methods tend to split traveling Gaussian waves into separate latent components, we used the percentage of explained variance of the first eigenvector as a quality metric. The greater the explained variance, the more effectively zero-lag methods can capture traveling wave motion in a single latent component. We found a linear decrease in explained variance moving from values beyond 0.2 to larger values of the traveling index (**Figure 1B**). Overall, we found that for moderate values of the traveling index (< 0.5), the explained variance of the first eigenvector is greater than 80%. This suggests that zero-lag methods effectively capture a large majority of the variance of a spatiotemporal pattern with moderate traveling wave behavior. For systems consisting predominantly of standing waves, or a moderate degree of traveling waves, we expect solutions of time-lag and zero-lag FC methods to converge.

### 2. Three Prominent Spatiotemporal Patterns

To understand the standing and traveling wave components that underlie empirical spontaneous BOLD fluctuations, we applied CPCA to a random sample (*n*=50) of resting-state fMRI scans (Human Connectome Project Young Adult; HCP: YA S1200). We downsampled the surface-based cortical time series to approximately 5,000 vertices. All analyses were successfully replicated in an independent sample (*n*=50) of unrelated HCP subjects (**Supplementary Figure 1** and **Supplementary Figure 2**).

To choose the number of CPCA components to extract, we utilized the Catell’s scree plot test applied in our simulations (**Figure 1A**). This criterion indicated a clear flattening in explained variance beyond three principal components (**Supplementary Figure 3**). The three leading complex principal components represent the top three dimensions of variability across complex-valued cortical BOLD time series. Associated with each complex principal component is a phase delay map, reflecting the time-delay (in seconds) between cortical vertices (see ‘Methods and Materials’). To examine the temporal progression of each complex principal component, we sampled the reconstructed BOLD time series from each component at multiple, equally-spaced phases of its cycle (*n_timepoints_*= 30; see ‘Methods and Materials’). Movies of the reconstructed BOLD time courses are displayed in **Supplementary Movie 1**. Detailed descriptions of the spatiotemporal patterns of each complex principal component, and comparisons of their propagation patterns, are provided in **Supplementary Figure 4**. Complex principal component phase delay maps beyond the first three are in **Supplementary Figure 5**.

The first component (*pattern one*), representing the leading axis of variance, explains 21.4% of the variance in intrinsic BOLD time series, greater than three times the variance explained by the second (6.8%) or third component (5.7 %). The traveling index of the first component was 0.25, indicating a largely standing spatiotemporal pattern, with some traveling wave behavior. The dynamics can be separated into two phases. In the first phase, negative cortex-wide BOLD amplitudes are observed that are strongest in the sensorimotor cortex (SM), superior parietal lobule (SP), lateral visual cortex (LV), and superior temporal (ST) gyrus (**Figure 2A; Movie 1**). We refer to these brain regions as the somato-motor-visual (SMLV) complex, noting that it also includes some regions outside sensory-motor cortices (e.g. SP and ST). The second phase exhibited a propagation of strong negative BOLD amplitudes in the SMLV toward cortical regions overlapping primarily with the frontoparietal network (FPN), but also with the DMN and V1. This entire spatiotemporal sequence of negative BOLD amplitudes was followed by a spatiotemporal sequence with positive BOLD amplitudes with the same dynamics.

**Figure 2.**
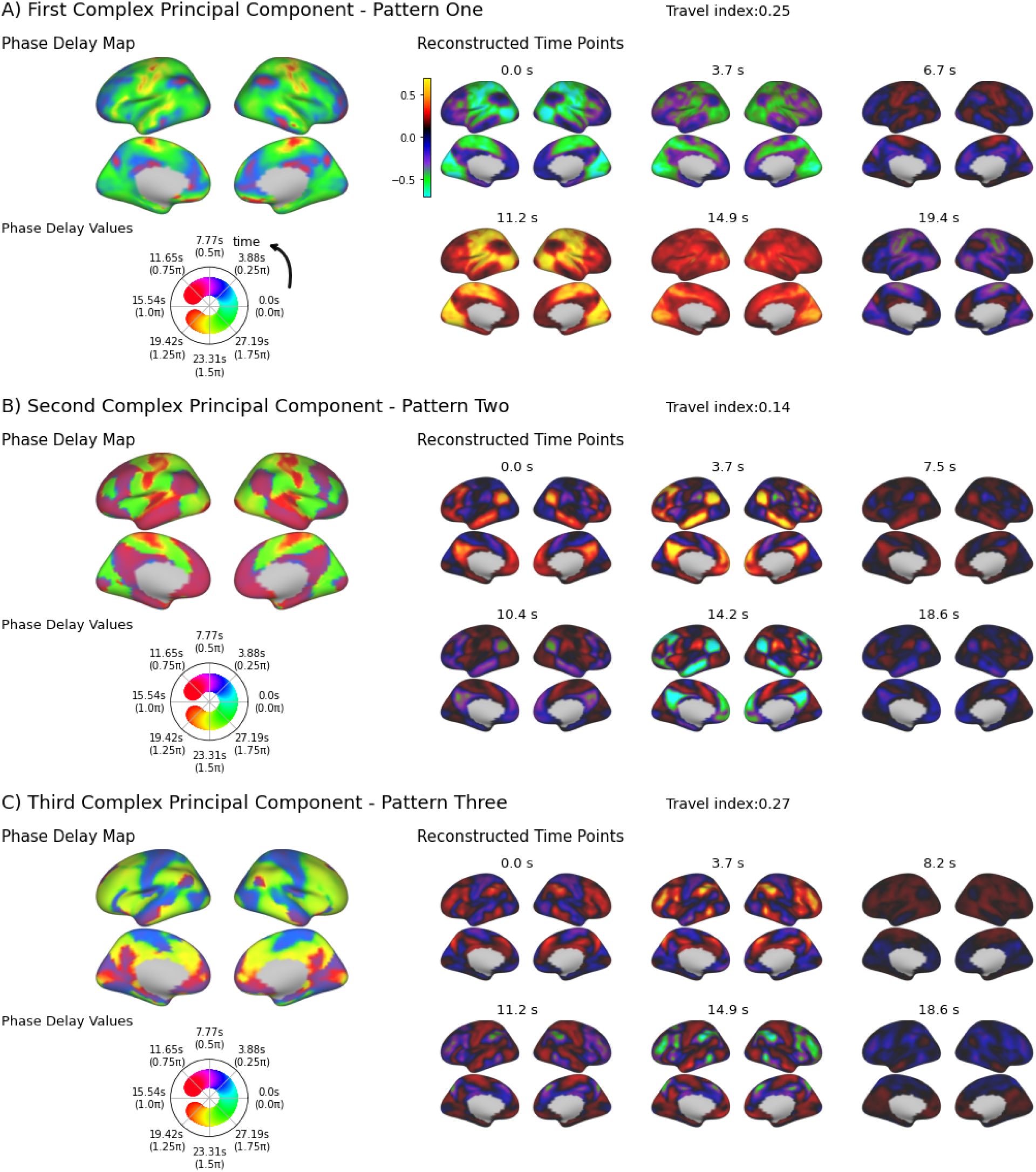
Form and Properties of Three Prominent Spatiotemporal Patterns. Phase delay maps and reconstructed time points of the first three complex principal components. Phase delay maps represent the temporal ordering (in seconds) of cortical vertex BOLD time series within the spatiotemporal pattern. Phase delay maps describe a repeating or cyclical pattern expressed in radians (0 to 2π) around a unit circle, where a phase value of 0 corresponds to the beginning of the spatiotemporal pattern, and 2π corresponds to the end of the spatiotemporal pattern. For clarity, radians are converted to temporal units (seconds) (see ‘Methods and Materials’). The values in the phase delay map correspond to the temporal delay (in seconds) between two cortical vertices, such that smaller values occur *before* larger values. Values are mapped to a cyclical color map to emphasize the cyclical temporal progression of each spatiotemporal pattern. To illustrate the temporal progression of the spatiotemporal patterns, six reconstructed time points are displayed for each pattern. The phase delay map and reconstructed time points are shown for **A**) the first spatiotemporal pattern - ‘pattern one’, **B**) the second spatiotemporal pattern - ‘pattern two’, and **C**) the third spatiotemporal pattern - ‘pattern three’.

The second component (*pattern two*) was the most stationary of the first three components, with a traveling index of 0.14. The overall spatiotemporal pattern can be described as an anti-correlated oscillation between SMLV regions and the DMN. Interestingly, this pattern of anti-correlation or bipolar contrast appeared to delineate the unipolar (all-negative) contrast in the first phase of the first complex principal component. Visual inspection revealed a minor traveling wave pattern propagating out from the SM region to the premotor cortex (anterior direction) and SP (posterior direction) in-between peak amplitudes of the anti-correlated oscillation.

The third component (*pattern three*) was similar to the first component with a traveling index of 0.27. The dynamics can be split into two phases. In the first phase, strong negative amplitudes are observed in the SM, ST and LV, with weak negative amplitudes in the DMN; strong positive amplitudes are observed in the inferior parietal lobule (IP), inferior temporal gyrus (IT), premotor cortex, dorsolateral prefrontal cortex (DLPFC) and V1. The second phase was marked by propagation from the IP (anterior direction) and premotor (posterior direction) towards the SM, and the IT (posterior direction) towards the LV.

To further understand what properties of the cortical BOLD time courses give rise to the time-lag structure of the three spatiotemporal patterns, we conducted two null model exercises. First, BOLD time courses were randomly shuffled to demonstrate that the time-lag structure of the spatiotemporal patterns depends on properties of the time courses beyond their zero-lag correlations. Random shuffling of the time courses preserves the zero-lag correlation structure, while eliminating time-lag correlation structure. As expected, CPCA of time point shuffled data removes the intermediate phase delays (between 0 and π), leaving behind a pure standing wave structure (**Supplementary Figure 6)**.

Second, we simulated time courses from a first-order multivariate autoregression (VAR(1)) modelfit to the cortical BOLD time series. This model leaves the time-lag (i.e. autoregressive) structure of the cortical BOLD time courses intact, while assuming Gaussianity, linearity and stationarity of the time courses ^23^. CPCA of the simulated time courses generated from the VAR(1) model replicated both the zero-lag and time-lag structure of the three spatiotemporal patterns (**Supplementary Figure 6**). These findings suggest that the three spatiotemporal patterns arise from properties of the BOLD time courses that are mostly Gaussian, linear and stationary.

### 3. FC Topographies Reflect the Three Spatiotemporal Patterns

The three spatiotemporal patterns are predominantly composed of standing waves. This finding, and the results of the simulation study (**Figure 2**), suggest that zero-lag FC methods should capture the majority of the variance in these spatiotemporal patterns. We selected a large number of commonly-used zero-lag FC methods and applied them to the original time courses from the same resting-state fMRI data. Dimension-reduction methods included principal component analysis (PCA), PCA with Varimax rotation (varimax) ^24^, Laplacian Eigenmaps (LE)^25^, spatial independent component analysis (SICA) ^26^, and temporal independent component analysis (TICA)^16^. Hidden Markov models (HMM), commonly used latent state space models for estimating brain states^17^, were also included. Seed-based FC analysis methods included seed-based correlation analysis^6^ and co-activation pattern (CAP) analysis ^15^ with k-means clustering of suprathreshold time points into two clusters. Seed-based methods were run for three key seed locations corresponding to major hubs in the SMLV, DMN , and FPN - the somatosensory cortex, precuneus, and dorsolateral prefrontal cortex^4^. Results were identical for alternatively-placed seed regions within these three networks (**Supplementary Figure 7**).

To determine a meaningful number of dimensions in our latent dimension-reduction analyses (PCA, varimax PCA, LE, SICA, TICA and HMM), we again used the scree plot criterion. The scree plot of the zero-lag correlation matrix indicated a clear flattening in explained variance after three principal components (**Figure 2C**).

To compare FC topography spatial similarity, we used the spatial correlation between the cortical vertex weight values of each topography. To summarize FC topography similarities, we compared each topography to the first three principal component (PC) maps computed from PCA. Similar to the first complex PC of CPCA, the first PC explains 20.4% of the variance in BOLD time series; greater than three times the variance explained by the second (6.8%) or third PC (6.1%). Each FC topography exhibits strong similarities with one or more of the first three PCs (***r* > 0.6**) (**Figure 3A**). Statistical significance of the spatial correlation between each FC topography and its most strongly correlated PC was computed using spin permutation tests ^27^ (N_samples_ = 1000). All correlation pairs were statistically significant (p = 0.001; lower limit of permutation test). However, there was notable variability in the strength of correlations between each FC topography and one or more of the PCs. Further, strong similarities between FC topographies are seen for more than one PC. Detailed results of seed-based regression/CAP and dimension-reduction analyses are provided in **Supplementary Figures 8** and **9**. Overall, our survey revealed a considerable consistency in FC topographies across methods.

**Figure 3.**
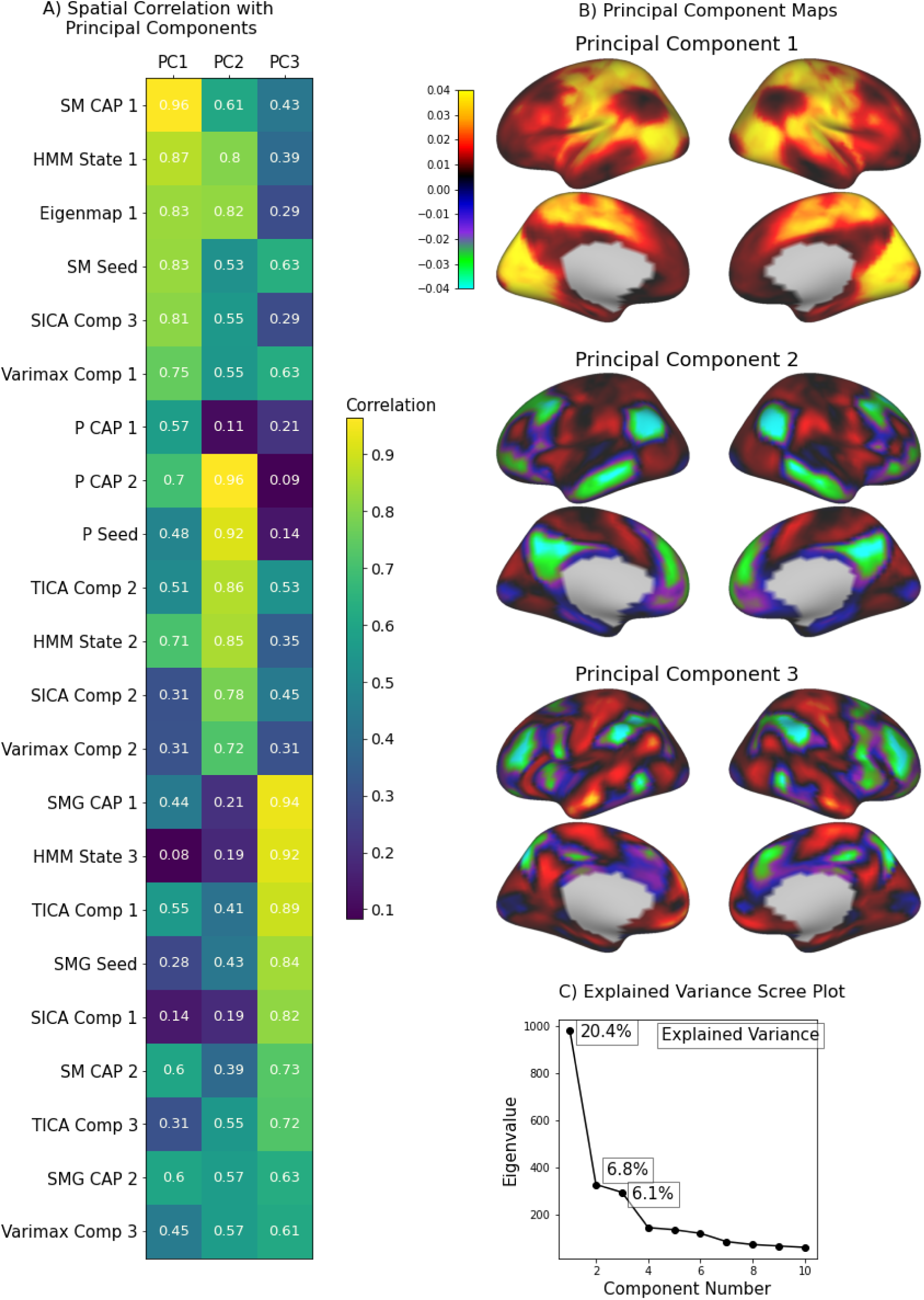
Form and Properties of Three Prominent Functional Connectivity Topographies. **A)** The spatial absolute-valued correlation between the first three principal component maps and each FC topography displayed as a table. All correlations are rounded to the second decimal place. The color of each cell in the table is shaded from light yellow (strong correlation) to dark blue (weak correlation). All FC topographies in our survey exhibited strong spatial correlations (Pearson’s correlation) with one (or two) of the first three principal components. **B)** The first three principal component spatial maps. **C)** The scree plot that displays the explained variance in cortical time series for each successive principal component. The scree plot indicates a clear elbow after the third principal component, indicating a ‘diminishing return’ in explained variance of extracting more components. (HMM = hidden markov model; TICA = temporal ICA; SICA = spatial ICA; P=precuneus; SM= somatosensory; SMG=supramarginal gyrus; Clus=cluster; Comp=component; PC = Principal Component).

Comparison of the spatial weights of the three PCs (**Figure 3B**) with the spatiotemporal patterns (patterns one, two and three) of the three complex PCs (**Figure 2**) illustrates that both methods capture similar spatial patterns. The correspondence was one-to-one; the first PC matching pattern one, the second PC matching pattern two, and the third PC matching pattern three. The static spatial weights of each PC appeared as a ‘snapshot’ within their corresponding spatiotemporal pattern: PC1-pattern one (***r* = 0.998**, t = 11.9s), PC2-pattern two (***r* = 0.986**, t = 12.6s), PC3-pattern three (***r* = 0.972**, t = 3.7s). Further, the three spatiotemporal pattern time courses closely tracked the first three principal component time courses: PC1-pattern one (***r* = 0.98**), PC2-pattern two (***r* = 0.95**), and PC3-pattern three (***r* = -0.83**; temporal lag of ∼3 TRs). Given the similarity, we refer to the patterns produced by standard PCA and CPCA, as pattern one, two and three, interchangeably.

### 4. Comparison With Previously Observed Resting-State Phenomena

A further aim of this study was to understand the relationship between these three spatiotemporal patterns and previously observed phenomena in intrinsic BOLD signals. Lag projections^7^ and the QPP^18^ correspond to time-lagged phenomena at shorter (∼2s) and longer (∼20s) time scales, respectively. Lag projections are computed from the average pairwise time-delays between BOLD time courses, and represent the average ‘ordering’ in time of BOLD amplitude peaks across the brain. An analogous time delay representation can be computed from the average of the pairwise phase delays of the complex correlation matrix used by CPCA. The spatial correlation between the lag projection and the CPCA-derived lag projection were very similar (***r* = 0.83**), and both exhibit the same direction of propagation (**Figure 4**). Interestingly, both average latency structures exhibited a strong spatial similarity with the phase delay map of pattern one (lag projection: ***r* = 0.81**; circular mean: ***r* = 0.98**) (**Figure 4**). This suggests the average latency structure of spontaneous BOLD fluctuations is largely driven by the first spatiotemporal pattern. Note, due to preprocessing differences the lag projection we computed only partially resembles the average lag projection in Mitra et al.^7^ - as discussed in **Supplementary Figure 10.**

**Figure 4.**
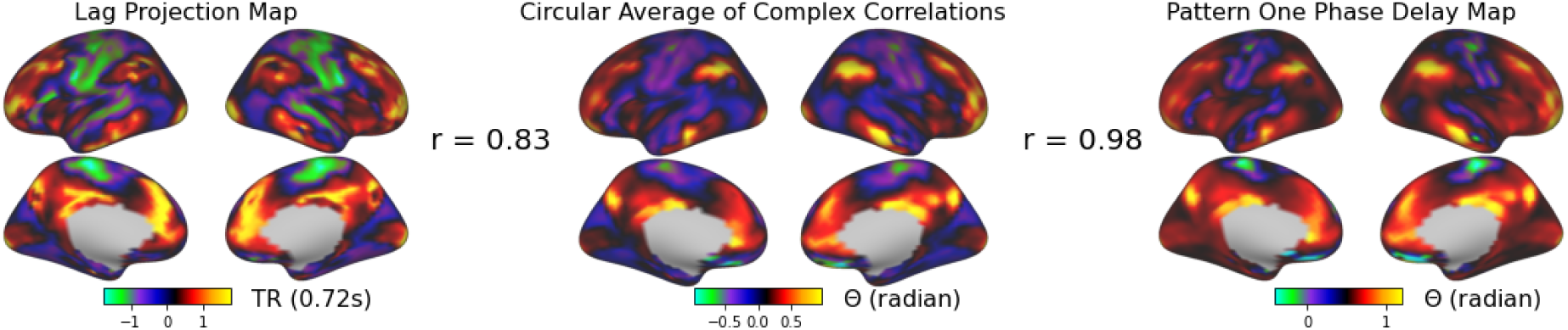
Similar Propagation Patterns between Average Latency Structure and Pattern One. Comparison between the average latency structure of spontaneous BOLD fluctuations and the phase delay map of pattern one. The lag projection map (left) represents the average time delay (in seconds) between each vertex of the cortex. The circular average map (middle) represents the average phase delay (in radians) between each vertex of the cortex. As in Figure 2, the pattern one phase delay map (right) represents the phase delay (in radians) between the vertices within the dynamics of pattern one. The two methods for computing average latency structure exhibiting strong agreement in their propagation patterns **(*r* = 0.83**). The strong similarity between the average latency structure (circular average) and pattern one phase delay map (***r* = 0.98**) indicates that the average latency structure is primarily driven by the spatiotemporal dynamics of pattern one.

The QPP is conventionally derived from BOLD time courses after the application of global signal regression. As shown by Yousefi et al. ^28^, two types of QPP can be observed across individuals: one QPP pattern exhibits a global pattern of activity with positive correlation across brain regions (‘global’ QPP), and the other QPP exhibits a global pattern of anti-correlation between the TP and DMN (‘anticorrelated’ QPP). Global signal regression was found to consistently produce the anti-correlated QPP across all individuals.

We hypothesized that the global QPP and anti-correlated QPP would correspond to patterns one and two, respectively. To derive the global QPP, we applied the QPP algorithm without global signal regression. To derive the anti-correlated QPP, we applied the QPP algorithm after global signal regression was applied to all BOLD time courses. As hypothesized, the time course of the global QPP derived from non-global signal regressed time courses was strongly correlated with the time course of pattern one (***r* = 0.71**) at a time-lag of 7TRs (∼5s). Further, the time course of the anti-correlated QPP was strongly correlated with pattern two (**r = 0.80**) at a time lag of 16 TRs (∼12s). We visualized the spatiotemporal template of the global and anti-correlated QPP from the repeated template-averaging procedure, and compared it to the time point reconstruction (see above) of pattern one and pattern two, respectively. The temporal dynamics of pattern one and the global QPP overlapped significantly (**Supplementary Movie 2)**, as did the temporal dynamics of pattern two and the anticorrelated QPP (**Supplementary Movie 3**).

At peak amplitudes of pattern one, the distribution of weights is roughly all positive or all negative (**Figure 2A**). This suggests that pattern one may track the global mean time course of cortical vertices, or the ‘global signal’. Indeed, the time course of pattern one and the global mean time course are statistically indistinguishable (***r* = 0.97**). This also suggests that the temporal dynamics surrounding the time points before and after the peak of the global signal correspond to the dynamics of pattern one. We constructed a dynamic visualization of the global signal through a peak-averaging procedure. Specifically, all BOLD time courses within a fixed window (15TRs on each side) were averaged around randomly-sampled peaks (N=200, > 1 standard deviation above the mean) of the global mean time course. This revealed that the temporal dynamics surrounding peaks of the global signal precisely match those of pattern one (**Supplementary Movie 2**).

The temporal dynamics of pattern two largely represent an anti-correlated pattern between the SMLV and the DMN. This resembles the task-positive/task-negative anti-correlation pattern originally observed by Fox et al.^6^ and Fransson^29^. We reproduced these results by creating whole-brain vertex-wide correlation maps using a seed time course from the left and right precuneus; a key DMN node. As expected, an anti-correlated pattern was observed between the SMLV and DMN (**Figure 5A**). Additionally, the precuneus-seed whole-brain correlation spatial map precisely corresponds to the pattern of BOLD activity at peak amplitudes of pattern two (CPC1: ***r* = 0.92**, t = 1.8s). This suggests that the task-positive vs. task-negative pattern arises from the anti-correlated dynamics between the SMLV and DMN represented by pattern two.

**Figure 5.**
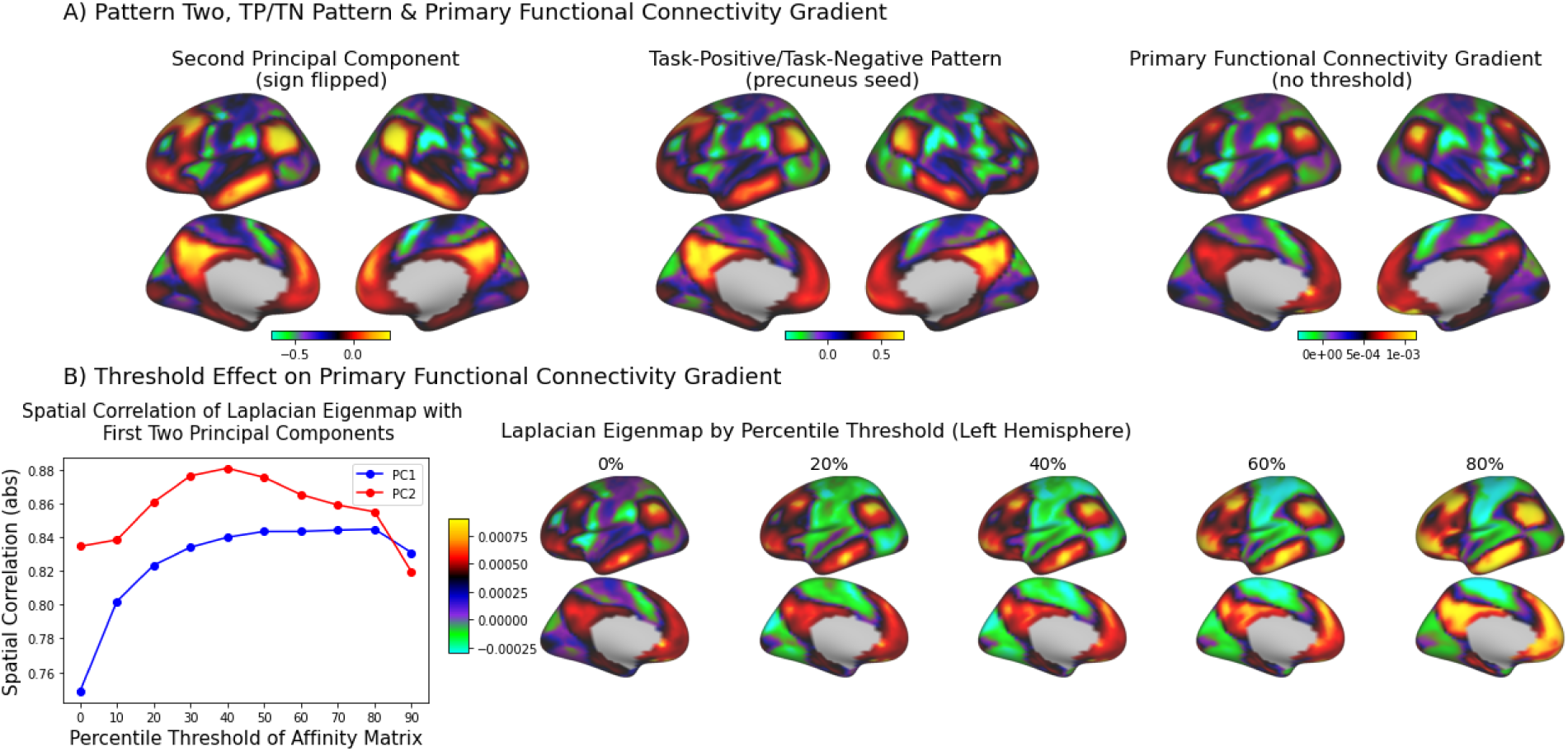
The Task-Positive/Task-Negative Pattern, Primary Functional Connectivity Gradient, and Pattern Two Describe the Same Spatiotemporal Pattern. **A**) From left to right, pattern two, task-positive/task-negative (TP/TN) pattern, and the PG represented by the spatial weights of the second principal component from PCA (sign flipped for consistency), seed-based correlation map (precuneus seed), and first Laplacian eigenmap with no thresholding of the affinity matrix, respectively. Similar spatial patterns are produced from all three analyses - pattern two: TP/TN (***r* =0.96**) and pattern two:PG (***r* = 0.83**). **B**) The effect of functional connectivity (FC) matrix percentile thresholding on the resulting spatial weights of the PG, computed as the first eigenmap of the Laplacian Eigenmap algorithm (left hemisphere). At zero to low-thresholding of the FC matrix, the first Laplacian Eigenmap resembles pattern two (PC2). As the threshold is raised, the spatial weights of vertices within the DMN and SMLV become more uniform. At higher thresholds this results in an Eigenmap that resembles pattern one.

A similar anti-correlation pattern to the task-positive/task-negative pattern has been observed in the FC gradient literature ^2^, and is known as the principal FC gradient (PG). In our zero-lag FC survey (**Figure 3**), we computed the PG as the first component derived from the Laplacian Eigenmaps (LE) algorithm, consistent with Vos de Wael et al.^25^ The PG resembles the anti-correlated spatial structure of pattern two, but this similarity depends on the level of thresholding of the FC matrix input to the LE algorithm (**Figure 5B**). The FC matrix represents Pearson’s correlation of BOLD time courses between all pairs of cortical vertices (i.e. correlation matrix). Generally, thresholding is performed row or column-wise on the FC matrix (e.g. 90th percentile of correlation values within that row) ^2^. If no FC matrix thresholding is performed, the first eigenmap produced from LE precisely resembles the BOLD activity in pattern two (PC2: ***r* = 0.83**; **Figure 5A**). As the percentile threshold applied to the FC matrix increases, the spatial weights of vertices within the DMN and SMLV become more uniform (**Figure 4B**). At higher percentile thresholds (e.g. 90th percentile), a contrast between the SMLV and DMN appears that is similar to the unipolar contrast of pattern one (PC1: ***r =* 0.83**) and the anti-correlation contrast of pattern two (PC2: ***r* = 0.82**). The reason that the first component of LE returns pattern two, over the more prominent pattern one, is due to time-point centering (**Supplementary Figure 11**).

### 5. Network-Based Representations of Functional Connectivity

The network or graph-based approach to FC analysis analyzes the structure of pairwise relationships between BOLD time courses. We investigated the degree to which the structure of pairwise relationships between BOLD time courses arises from the dynamics of the three spatiotemporal patterns. A FC matrix was constructed by computing correlations between all pairs of cortical BOLD time courses (**Figure 6**). We compared this matrix to the reconstructed FC matrix from the three spatiotemporal patterns. We estimated the similarity between the two FC matrices by computing the correlation coefficient between the lower triangles of each matrix. Despite a larger mean correlation in the reconstructed FC matrix, we found that the FC matrices were strongly correlated (***r* = 0.77**).

**Figure 6.**
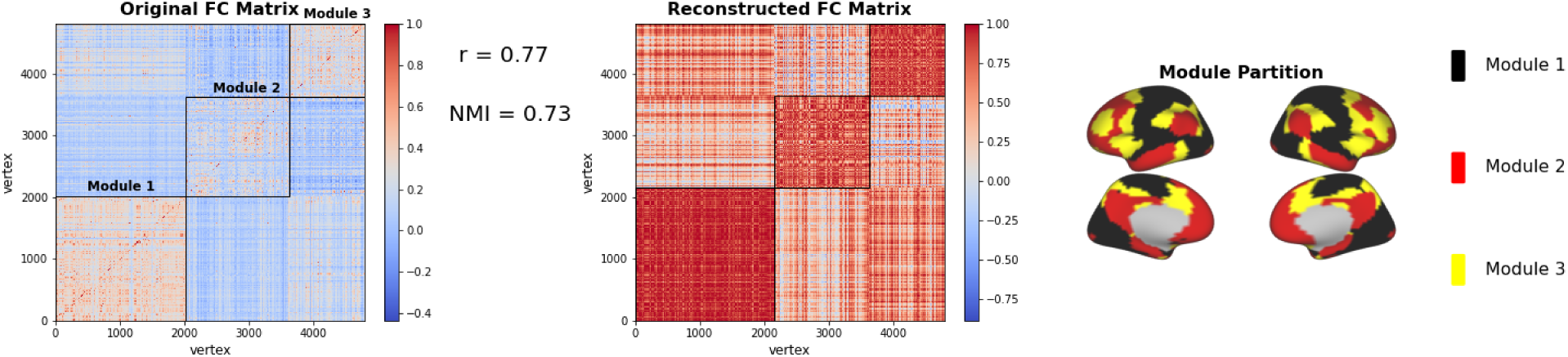
The Network Structure of Functional Connectivity Is Explained by the Three Spatiotemporal Patterns. Comparison of the correlation matrix of cortical BOLD time courses (left) with the correlation matrix of reconstructed cortical BOLD time courses (right) derived from the three spatiotemporal patterns, and the module assignments of each vertex (bottom). The rows and columns of the original and reconstructed correlation matrix are sorted and outlined (in black) according to the modular structure estimated from the Louvain modularity algorithm. The algorithm identified three primary modules in the SMLV, DMN and FPN. Despite a higher mean value of correlations in the reconstructed correlation matrix, the pattern of correlations between the two correlation matrices is highly similar **(*r* = 0.77**). Further, the modular structure of the original correlation matrix exhibits a high degree of similarity with the modular structure of the reconstructed correlation matrix (NMI = 0.73).

We also sought to determine whether the community structure of cortical BOLD time courses can arise from the shared dynamics of the three spatiotemporal patterns. We estimated network communities from the original and reconstructed FC matrix using the Louvain modularity-maximization algorithm. A measure of similarity between community structure, normalized mutual information (NMI), showed strong community structure similarity between the original and reconstructed FC matrix (NMI = 0.73).

## Discussion

The goal of this study was to provide a parsimonious taxonomy of prominent global spatiotemporal patterns in spontaneous BOLD activity to enable insight into the functional architecture of the human brain. Using a complex-valued extension of PCA, we identified three prominent spatiotemporal patterns consisting primarily of standing waves with some traveling wave behavior. Consistent with our simulations, the relative predominance of standing over traveling wave structure ensured that zero-lag FC methods effectively captured much of their spatiotemporal structure. This finding explains previous observations that traveling waves of BOLD activity resemble patterns of large-scale FC topographies ^8,9^.

An important finding from this study was the ubiquitous presence of three spatiotemporal patterns across a wide variety of zero-lag FC techniques. Several zero-lag FC methods (ICA, Varimax, HMM) involved an initial PCA dimension-reduction step. Thus, the common patterns resulting from zero-lag FC methods may be caused by a common reliance on PCA. However, an ample number of studies demonstrate that the output similarity across these methods is not guaranteed, but depends on the area of application ^13,30^ . Further, the same patterns emerged from zero-lag FC methods that do not rely on PCA (e.g., CAP and seed-based correlation) and also from time-lag analyses with no relation to PCA (QPP and lag projection).

Another core finding of our study is the identification of the brain’s average latency structure (**Figure 4**) with pattern one. This latency structure, originally discovered by averaging the pairwise time delays between brain regions^7^, was found to map directly to the phase delay pattern of pattern one. In other words, the average latency structure of spontaneous BOLD fluctuations reflects the traveling wave structure of the dominant spatiotemporal pattern (pattern one). Further, a repeated peak-averaging procedure demonstrated that this time-lag structure corresponds to local propagation patterns surrounding peaks of the global mean time course. This is consistent with previous findings of a global propagating wave event surrounding peaks of the global mean time course ^10^ .

While time-varying or dynamic FC was not examined in this study, fluctuations between the spatiotemporal patterns identified here may explain the variability in zero-lag FC structure over time. For example, a new time-varying FC analysis, edge time series analysis^31^, has found patterns of co-fluctuations in zero-lag FC resemble the spatial structure of the three spatiotemporal patterns^32^. While non-stationarity of zero-lag FC is a controversial topic ^33^, modeling work has shown that time-varying FC is consistent with stationary time series models^23^. Our null modeling results (**Supplementary Figure 6**) suggest that a stationary, linear auto-regressive process sufficiently describes the zero-lag and time-lag correlation structure of the three spatiotemporal patterns.

As the primary aim of our study was descriptive, we have avoided any explanatory or causal explanation of the possible neuronal or non-neuronal origins of these three spatiotemporal patterns. However, the identification of pattern one with the global mean time course suggests potential candidate causal mechanisms. One promising candidate is a systemic circulatory origin ^34^ . A significant portion of low-frequency BOLD signal variance is correlated with systemic oxygenation fluctuations in the periphery ^11^. This low-frequency peripheral oscillation tracks the global mean fMRI time course^35^ and exhibits significant traveling wave behavior induced by differential blood transit time in the cerebral vasculature ^36^. Potential sources of this systemic circulatory effect include vasomotion, fluctuations in arterial CO2 and/or Mayer waves ^34^. However, other studies support a neural origin for the global signal ^37,38^. We recently demonstrated that individual differences in global signal topography in the human brain are related to life outcomes and psychological function ^19^. This work follows studies using pharmacological intervention^38^ and electrophysiological recordings in macaques ^37^ providing evidence for neural origins of the global signal. Momentary fluctuations in arousal and/or vigilance are also related to the global BOLD signal ^37^. Some physiological processes strongly co-vary with cortical excitability ^39^, making it even more difficult to disentangle the neural vs. non-neural sources of the BOLD fluctuations. Taken together, these findings suggest that the global signal (and by extension pattern one), while influenced by vascular and other systemic physiological sources, is at least partially driven by neuronal sources. The current study does not resolve these questions about the sources of the BOLD fluctuations, and underscores the importance of understanding vascular components of the global signal in order to effectively denoise fMRI data while preserving neuronal signals ^40^.

Candidate origins of patterns two and three are more difficult to identify. Pattern two closely resembles the task-negative/task-positive pattern originally discovered in resting-state fMRI^6^. A similar anti-correlated pattern is consistently observed in response to task stimuli ^41^. Pattern two strongly resembles the QPP obtained after global signal regression, which has been linked to infraslow electrical activity ^42,43^. Patterns of propagating activity have also been observed in optical fluorescence imaging ^44^ and electrophysiological recordings^45^, which demonstrates that time-lagged relationships can arise from neural as well as hemodynamic processes. Future research may be directed towards a more complete understanding of the common or distinct neuronal or physiological mechanisms that give rise to these spatiotemporal patterns.

While these three spatiotemporal patterns explain a significant portion of variance in spontaneous BOLD fluctuations (∼32%), there is a considerable portion of variance left unexplained. Notably, these three components did account for much of the observed whole-brain functional connectivity. Thus, the remaining, larger fraction of variance, may contribute less to the global spatial structure of BOLD signals. The sources of this remaining signal component remain unclear. While some portion is likely to arise from non-neural effects, there is an intriguing possibility that it also contains activity related to ongoing cognition^46^. We speculate that it may reflect more spatially focal variance in spontaneous BOLD fluctuations, possibly more closely related to neurovascular coupling than the global patterns observed in this study.

It is important to qualify the assumptions on which our comparisons were based. First, the primary metric that determined the number of latent dimensions in our analysis was explained variance. The number three was chosen based on the observation of diminishing explained variance with increasingly higher-dimensional solutions. While alternative methods for the choice of dimensionality have been proposed, we found that the criterion of explained variance achieved high robustness. Second, most of the analysis approaches we surveyed can reveal finer-grained spatial insights at higher component or cluster numbers (e.g., ICA or Varimax-PCA). However, we do not expect the same consistency in analytic approaches at finer-grained levels of analysis. Empirical examination of consistency in zero-lag FC analyses at solutions greater than three components confirms this: higher dimensional solutions yield less consistent FC topographies (**Supplementary Figure 12**). Importantly, the consistency of analysis approaches at our low-dimensional level of analysis suggests that these three spatiotemporal patterns are robust phenomena. Finally, these three spatiotemporal patterns were modeled at the population level; intersubject variability in the spatial structure of these patterns was not considered in our analyses. However, an examination of the first three principal components across a sample of subjects reveals notable intersubject variability in their spatial structure (**Supplementary Figure 13**).

In summary, we identified three spatiotemporal patterns that parsimoniously recapitulate major findings from a wide range of analytical techniques. Further, these patterns account for much of the large-scale structure that underlies all FC analysis, and therefore impact how we interpret everything from functional networks to graphs. As the study of the brain as a complex system advances, these three spatiotemporal patterns have potential applications in better constraining generative models of brain activity, predicting variability in response to external stimuli, informing targeted modulation of brain activity to achieve a particular state, and understanding interactions between global brain states and localized activity in neuronal circuits. The concise characterization of systems-level brain activity in three spatiotemporal patterns will facilitate the cross-scale research needed to link fundamental neuroscience studies and human behavior.

## Supplementary Materials

### Supplementary Movies

*Supplementary Movie 1 - Visualization of Three Spatiotemporal Patterns*

Supplementary Movie 1. **Visualization of Three Spatiotemporal Patterns.** Temporal reconstruction of all three spatiotemporal patterns displayed as movies in the following order - pattern one, pattern two, and pattern three. The time points are equally-spaced samples (N=30) of the spatiotemporal patterns. The seconds since the beginning of the spatiotemporal pattern are displayed in the top left. In the bottom of the panel, the time points of the spatiotemporal pattern are displayed in three-dimensional principal component space (**Figure 2**). Two-dimensional slices of the three principal component space (see **Figure 3**) are displayed as the three 2-dimensional plots. The progression of time points in the principal component space is illustrated by a cyclical color map (light to dark to light). The movement of the spatiotemporal pattern through this space is illustrated by a moving red dot from time point-to-time point in synchronization with the temporal reconstruction in the movie.

*Supplementary Movie 2 - Dynamic Visualization of the Quasiperiodic Pattern, Pattern One, and Global Signal*.

Supplementary Movie 2. **Dynamic Visualization of the Quasiperiodic Pattern, Pattern One, and Global Signal.** The 30 time points (TR=0.72s) of the global QPP, pattern one, and peak-average global signal displayed as a movie (in that order). The time index of each sequence is displayed in the top left. The time points of pattern one are equally-spaced phase samples (N=30) of the time point reconstruction (see above). The time points of the global QPP are derived from the spatiotemporal template computed from the repeated-template-averaging procedure on non-global signal regressed data. The global signal visualization concatenates the left and right windows (w=15TRs) of the global signal peak-average. The time points of the global signal visualization begin at TR=-15, corresponding to 15 TRs pre-peak, and proceed to TR=15, corresponding to 15TRs post-peak.

*Supplementary Movie 3 - Anti-Correlated QPP and Pattern Two Visualization*

Supplementary Movie 3. **Dynamic Visualization of the Anti-Correlated Quasiperiodic Pattern and Pattern Two.** The 30 time points (TR=0.72s) of the anti-correlated QPP and pattern two. The time index of each sequence is displayed in the top left. The time points of pattern two are equally-spaced phase samples (N=30) of the time point reconstruction (see ‘Methods and Materials’). The time points of the anti-correlated QPP are derived from the spatiotemporal template computed from the repeated-template-averaging procedure on global signal regressed data.

### Supplementary Modeling Note - CPCA on Simulated Data

#### Simulation Approach and Methodology

To understand the behavior of zero-lag FC analyses in the presence of traveling wave oscillations, we conducted a simulation of traveling Gaussian waves on a two-dimensional grid. The grid was constructed as a 100-by-50 two-dimensional array of pixels. To operationalize the notion of standing and traveling waves we use the mathematical formalization of complex sinusoidal wave motion developed in Feeny^14^. A complex wave motion can be represented by the following function:

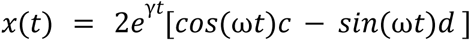

where *t* is the discrete-sampled time array, γ is an exponential decay term, ω is the angular frequency, and ***c*** and ***d*** are two vectors representing the two modes of the oscillation. The output of the function, ***x***, is a vector formed from the mixture of the two modes, ***c*** and ***d,*** weighted by a cosine and sine term (modulated by time, *t*). At the peak of the cosine oscillation, the sine oscillation is at zero, and the pattern encoded in vector c is the dominant configuration. At the peak of the sine oscillation, the cosine oscillation is at zero, and the pattern encoded in vector d is the dominant configuration. As a function of time, the output appears visually as cyclical ‘wave’ motion between the patterns defined by vectors *c* and *d*. These patterns can take any form or shape. For our simulation, the vectors represented flattened (i.e. converted to one-dimension) two-dimensional Gaussian curves with varying peak location. More specifically, ***d*** is a two-dimensional Gaussian curve (σ = 1) with its peak at one position in the 100-by-50 grid, and ***c*** is another two-dimensional Gaussian curve (σ = 1) with its peak lower down the 100-by-50 grid in the vertical direction. The simulation appears as a single two-dimensional Gaussian wave traveling from the Gaussian represented in vector ***c*** upwards to the Gaussian represented in vector *d*. The degree of traveling wave behavior of the complex motion is determined by the degree of statistical independence between the two Gaussian curve patterns. For example, for completely overlapping Gaussian curves (i.e. identical peak locations), the patterns are entirely dependent, which will appear as a pure standing oscillation. For completely non-overlapping Gaussian curves, the patterns are entirely independent, which will appear as a pure traveling wave oscillation.

To assess the behavior of zero-lag FC methods applied to oscillations with arbitrary mixtures of standing and traveling waves, we systematically varied the distance between the two Gaussian curves (vectors ***c*** and ***d***) by moving the peak location of the starting Gaussian curve (vector ***c*)** from the bottom of the 100-by-50 grid (no overlap) to the top of the grid (complete overlap with vector ***d***). In **Figure 1A**, we displayed the output of zero-lag FC analysis and CPCA at large (vertical peak distance of 50 pixels), moderate (peak distance of 25 pixels) and zero distances between the peaks of the two Gaussian curves. As noted above, large peak distance corresponds to approximately statistically independent Gaussian curve patterns (vectors ***c*** and ***d***), producing a pure traveling Gaussian wave. Zero peak distance corresponds to complete statistically dependent Gaussian curve patterns, producing a pure standing Gaussian wave. A simulated oscillation is run for each peak distance with the following parameters (ω = 1, γ = -1, number of cycles = 40, number of samples = 800). For each simulation, a limited amount of Gaussian noise (variance = 0.2) was added to each pixel time course. Movies files for the simulations display in **Figure 1A** are provided at the following link ( https://drive.google.com/drive/folders/1TWHY9Ktue7Y1mB6ITTYTf8FtCgqeDd05?usp=sharing).

#### I) Simulation of Two-Mode Traveling Wave Oscillations

The simulation illustrated in **Figure 1** consisted of a single traveling wave oscillation of two gaussian curves. To assess the ability of CPCA to separate multiple, overlapping traveling wave oscillations, we slightly modified the simulation presented in **Figure 1** (see ‘Methods and Materials’). In addition to the existing traveling wave oscillation between the two gaussian curves on the two-dimensional grid, we embedded a higher-frequency (ω = 2; see ‘Methods and Materials’) traveling wave oscillation of two smaller gaussian curves at identical peak locations. For this simulation, the two gaussian curves of each oscillation were completely separated, or statistically independent, thereby creating a pure traveling wave between the gaussian curves of each simulation. Identical parameters to simulation in the main text were used, excluding the angular frequency (ω; see ‘Methods and Materials’). This two mode simulation appeared as overlapping traveling wave oscillations between two lower frequency, large gaussian curves, and two higher frequency, small gaussian curves (**Supplementary Model Figure 1**). We applied CPCA to the pixel time courses to assess its ability to separate these oscillations into two latent factors.

**Supplementary Model Figure 1.**
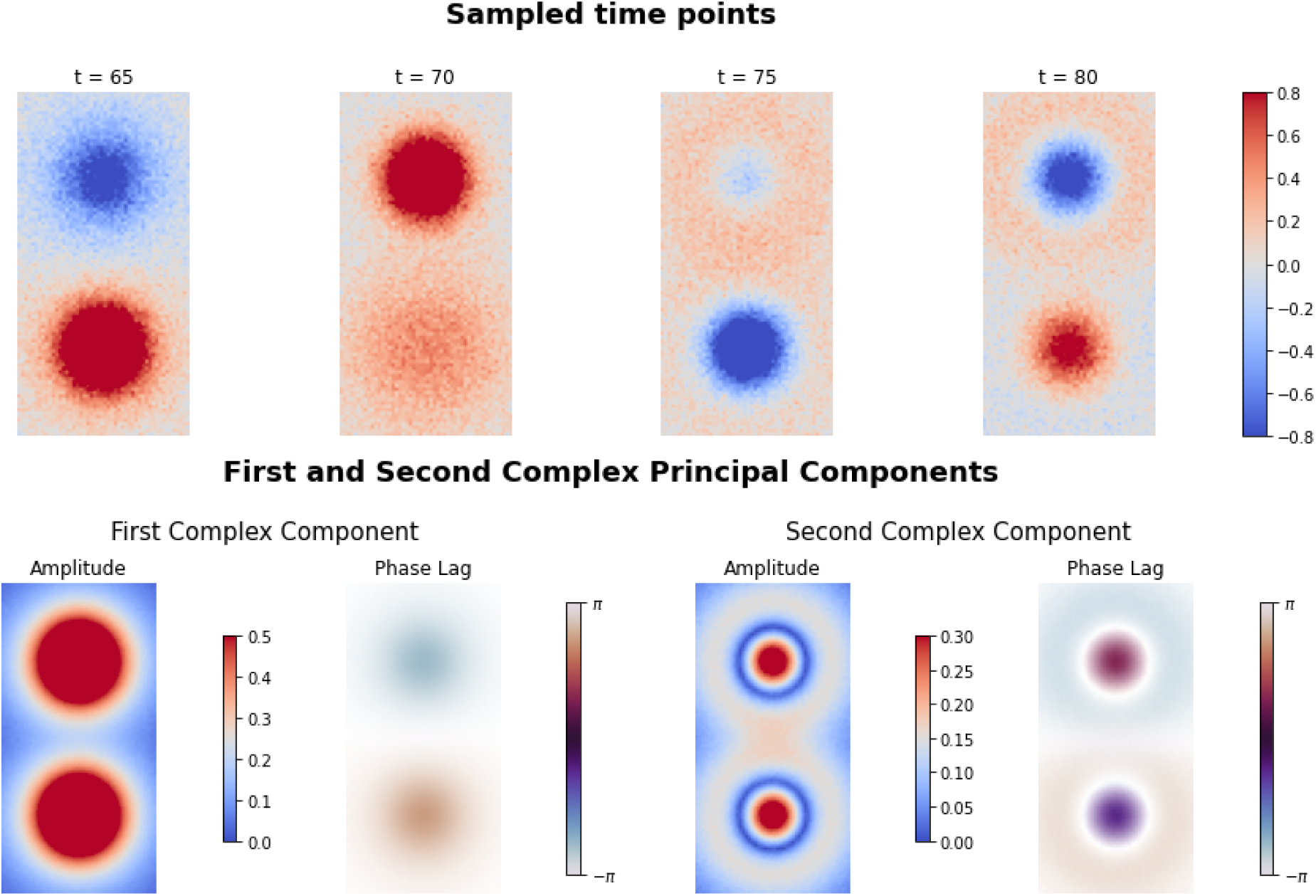
Simulation and Analysis of Two-Mode Traveling Wave Oscillations. Simulation of two spatially overlapping, traveling wave oscillations. The simulation consists of a high frequency oscillation between two smaller gaussian curves, and a low frequency oscillation between larger gaussian curves. Four sampled time points are displayed in the top panel. The amplitude and phase delay maps of the first two complex principal components estimated by CPCA are displayed in the bottom panel. As can be observed from the amplitude and phase delay map, CPCA accurately separated the two traveling wave oscillations into separate components.

We extracted two complex principal components from the complex-valued pixel time courses. We plotted the amplitude and phase lag of each component, representing the strength of ‘participation’ and time-lag of the pixels in the complex component, respectively. As indicated by the amplitude maps, the spatial outline of the larger and smaller gaussian curves is accurately separated into the first and second complex principal components, respectively. Further, the phase maps of the two complex principal components accurately distinguish the higher frequency oscillations of the smaller gaussian curves from the lower frequency oscillations of the larger gaussian curves.

#### II) Simulation of Traveling Wave Hemodynamic Signals

To demonstrate the ability of complex principal component analysis (CPCA) to extract spatiotemporal patterns from more complex signals, we applied CPCA to propagating fields of signals convolved with a hemodynamic response function. Impulse time series (1 for activation, 0 otherwise) convolved with the canonical hemodynamic response were spatially arranged along a square grid. The time series were arranged such that the time series in the top part of the square grid peaks early, and peaks later and later (time steps of 0.1 secs per row) as one moves down the grid. This arrangement provided a simple illustration of a global propagation event, where activity in one location travels to all other locations in a spatially continuous fashion. Gaussian noise was added to every time point of each time series, and slight phase and amplitude jitter are applied to each time series within a row of the square grid drawn from a uniform distribution. The global propagation event in the simulation was re-run 1000 times and temporally concatenated. It is important to note that this simulation was not intended to be a biologically realistic simulation of the mechanisms that produce observed BOLD propagation patterns. While this simulation is designed to have superficial similarities to spatial and temporal properties of BOLD propagation, we did intend to simulate its underlying data generating process. Below we display the first 20 time points of the global BOLD propagation simulation (**Supplementary Model Figure 2**.

**Supplementary Model Figure 2.**
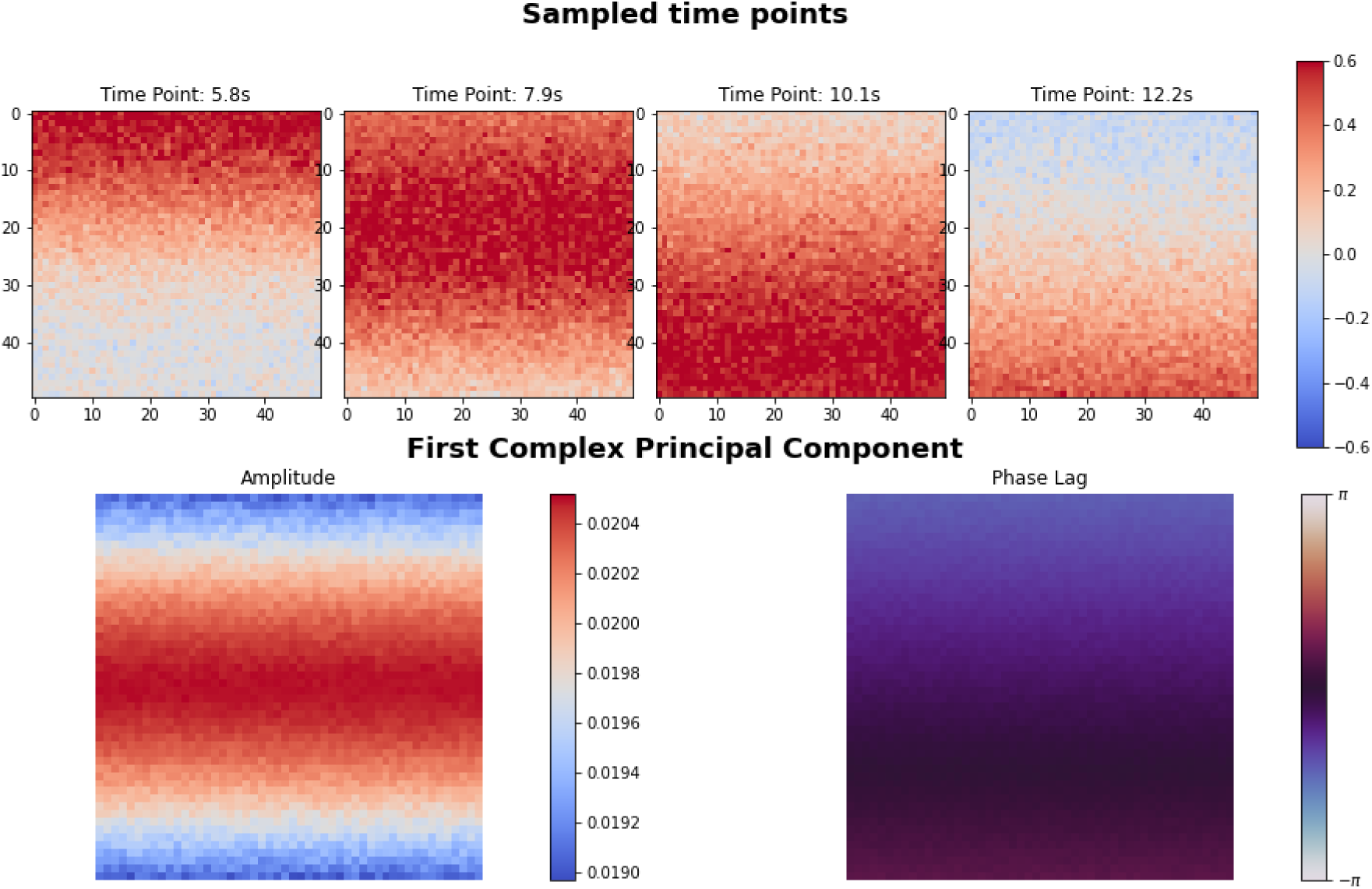
Global BOLD Propagation Simulation and Analysis. **Top panel:** Four sampled time points of an artificially constructed BOLD propagation simulation. Artificial ‘vertices’ are arranged along a 50-by-50 spatial grid. Vertex time series are created by convolving simple impulse time series with the canonical hemodynamic response function. The time-series of each vertex in the grid are time-lagged such that vertex time series in the upper part of the grid peak first, while those in the bottom part peak last. **Bottom Panel:** The amplitude and phase delay weights of the first complex principal component.

The sampled time points of the simulation (**Supplementary Model Figure 2**) illustrate the simulated global propagation event: peak BOLD amplitudes are first observed in the top of the square grid, followed by a subsequent propagation of peak BOLD amplitudes down the grid. Following the globally positive BOLD propagation event, there is a mirrored negative BOLD propagation event due to the post-response undershoot of the canonical hemodynamic response function. We extracted the first complex principal component from the complex time courses of the simulation. As illustrated in **Supplementary Model Figure 2**, the first complex principal component from CPCA accurately recovers the spatiotemporal pattern of the global propagation simulation. The phase delay map accurately describes the spatiotemporal pattern as BOLD activity that travels at a steady rate down the grid.

## Supplementary Figures

**Supplementary Figure 1.**
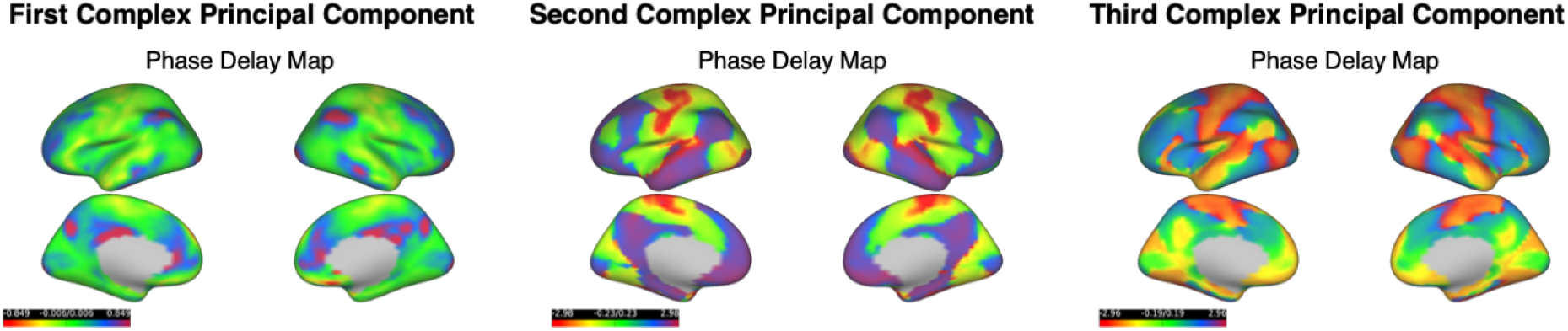
Replication of CPCA Phase Delay Maps in Independent Sample. **Phase Delay Maps of Complex Principal Components in an Independent Sample.** To give a sense of the robustness of our findings in an independent sample (unrelated to any subject in the primary sample), we display the CPCA phase delay maps derived from the independent sample. Displayed are the phase delay maps of the first three complex principal components in the independent replication sample of 50 subjects. Phase delay values are displayed in radians.

**Supplementary Figure 2.**
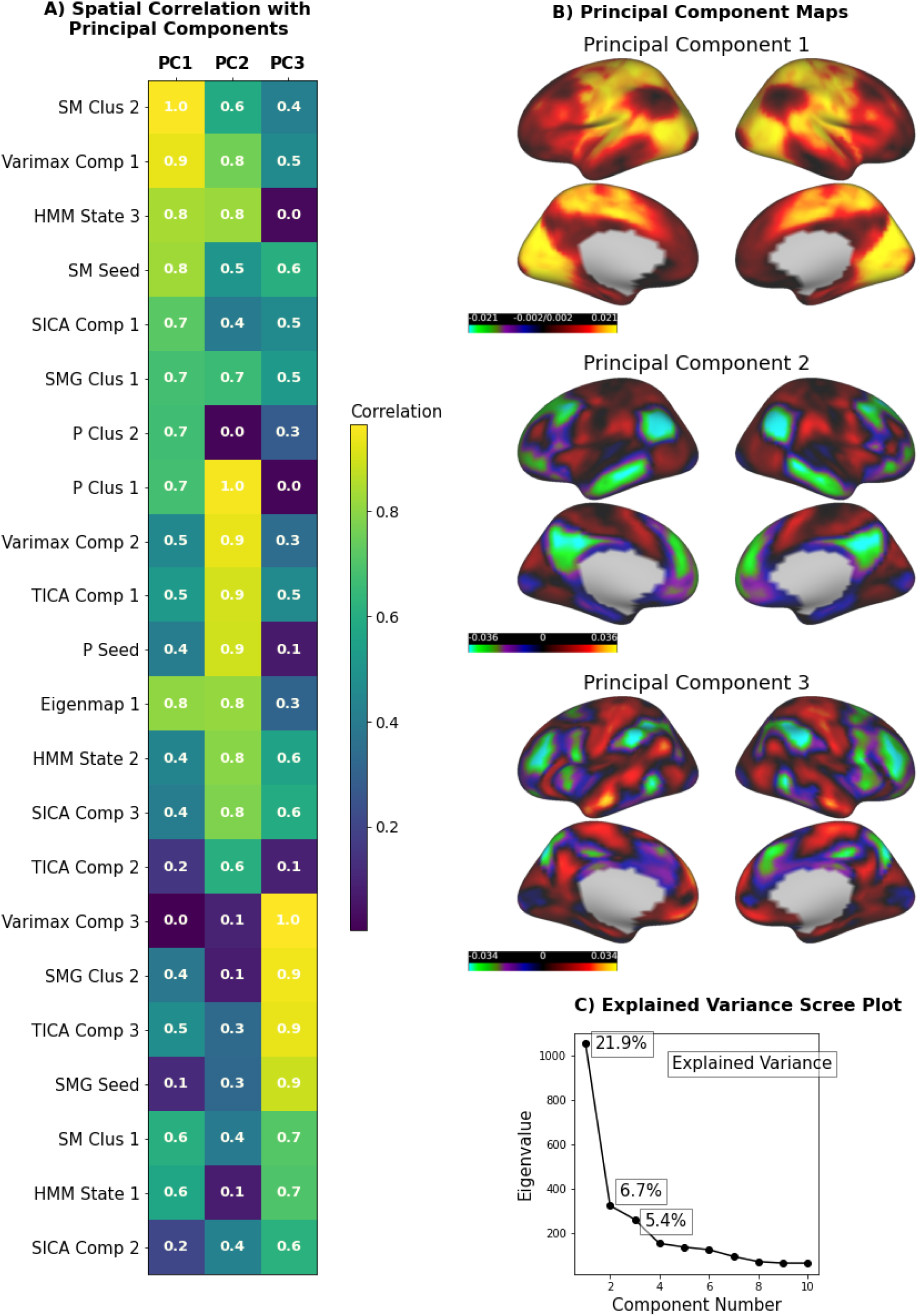
Replication of Zero-lag FC Survey in Independent Sample. **Zero-lag FC Survey in Independent Sample.** To give a sense of the robustness of our findings in an independent sample, we display zero-lag FC survey results (Figure 3) for the independent sample of 50 Subjects. **A)** The spatial absolute-valued correlation between the first three principal component maps and each FC topography displayed as a table. All correlations are rounded to the second decimal place. The color of each cell in the table is shaded from light yellow (strong correlation) to dark blue (weak correlation). **B)** The first three principal component spatial maps. **C)** The scree plot that displays the explained variance in cortical time series for each successive principal component.

**Supplementary Figure 3.**
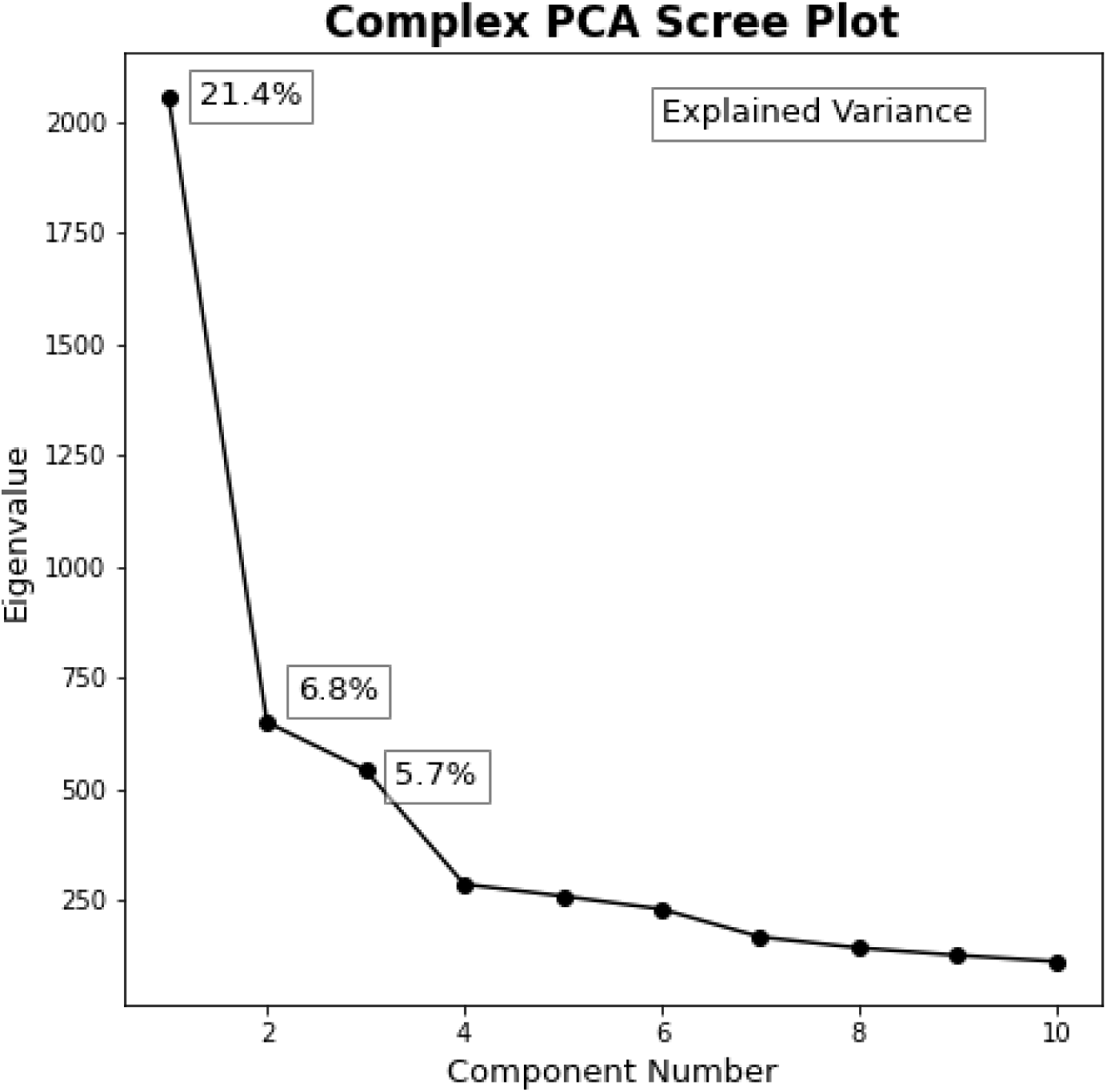
Complex Principal Component Scree Plot. **Scree Plot from Complex Principal Component Analysis.** The eigenvalue by component number plot (i.e. scree plot) used to determine the number of components to extract. There are clear elbows in the plot after one and three components, indicating a preferred solution of one or three principal components (three were chosen).

**Supplementary Figure 4.**
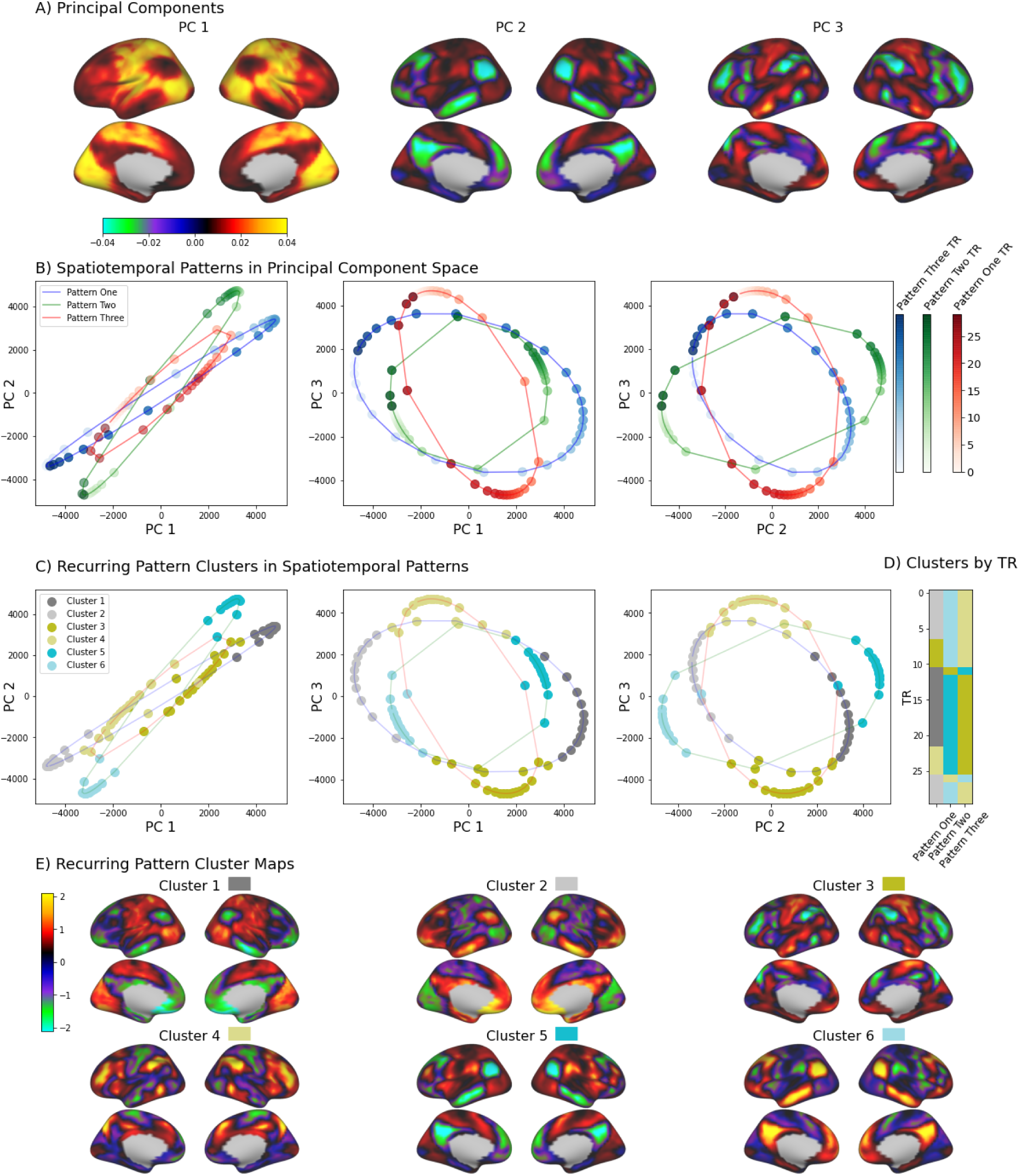
Steady States and Propagation Events in Spatiotemporal Patterns. **Spatiotemporal Patterns Consist of Steady States and Propagation Events That Repeat Across Patterns.** (PC = Principal Component) **A)** The spatial weights of the first three principal components. **B)** To visualize the temporal dynamics of the three spatiotemporal patterns (patterns one, two and three), we projected their reconstructed time points (see ‘Methods and Materials’) into the 3-dimensional space formed by the first three principal components. The time points are displayed on two-dimensional slices of each spatiotemporal pattern in the 3-dimensional principal component space - PC1-PC2, PC1-PC3, and PC2-PC3 spaces. The time points of patterns one, two and three are displayed as blue, green and red points, respectively. Consecutive time points of each spatiotemporal pattern are linked by lines. The time points of each spatiotemporal pattern are colored from light to darker to visualize the progression of time (N=30). The temporal cycle of each spatiotemporal pattern forms an oval in the three-dimensional principal component space, corresponding to a full cycle of the spatiotemporal pattern. For all three spatiotemporal patterns, most consecutive time points cluster closely together, indicating a ‘steady state’ of BOLD activity with relative stability of BOLD activity over that period. The steady states of pattern one, pattern two and pattern three vary strongest along the first, second and third principal component axes, respectively. These steady state periods are interrupted by large movement between consecutive time points that correspond to rapid propagation of BOLD activity towards another steady state. **C)** To examine repeating spatial patterns across the three spatiotemporal patterns, we applied a clustering algorithm (K-Means) to the reconstructed time points from all three spatiotemporal patterns. To avoid scaling differences in the distance calculations between time points, the BOLD values within each time point were z-score normalized. The same two-dimensional slices of each spatiotemporal pattern in the 3-dimensional principal component space are colored according to their cluster assignment by a k-means clustering algorithm. K-means clustering was used to identify recurring spatial patterns of BOLD activity across time points of the three spatiotemporal patterns. Six clusters were estimated. **D)** The cluster assignments (color) by time (*y-axis*) of each spatiotemporal pattern (*x-axis*). Note, that the same cluster assignment can occur across more than one spatiotemporal pattern. **E)** The cluster centroids from the k-means clustering algorithm, corresponding to the average spatial pattern of BOLD activity for the time points that belong to that cluster. Note, the cluster centroids of the first two clusters are mean-centered versions of the original unimodal (all-positive or all-negative) steady-state of pattern one, as z-score normalization of the time-points across vertices was performed beforehand. As revealed by the cluster solution, the six clusters correspond to the three pairs of mirrored or sign-flipped steady-states of the three spatiotemporal patterns. The first two clusters correspond to the steady states of pattern one. Clusters three and four correspond to the steady states of pattern three, and clusters five and six correspond to the steady states of pattern two.

**Supplementary Figure 5.**
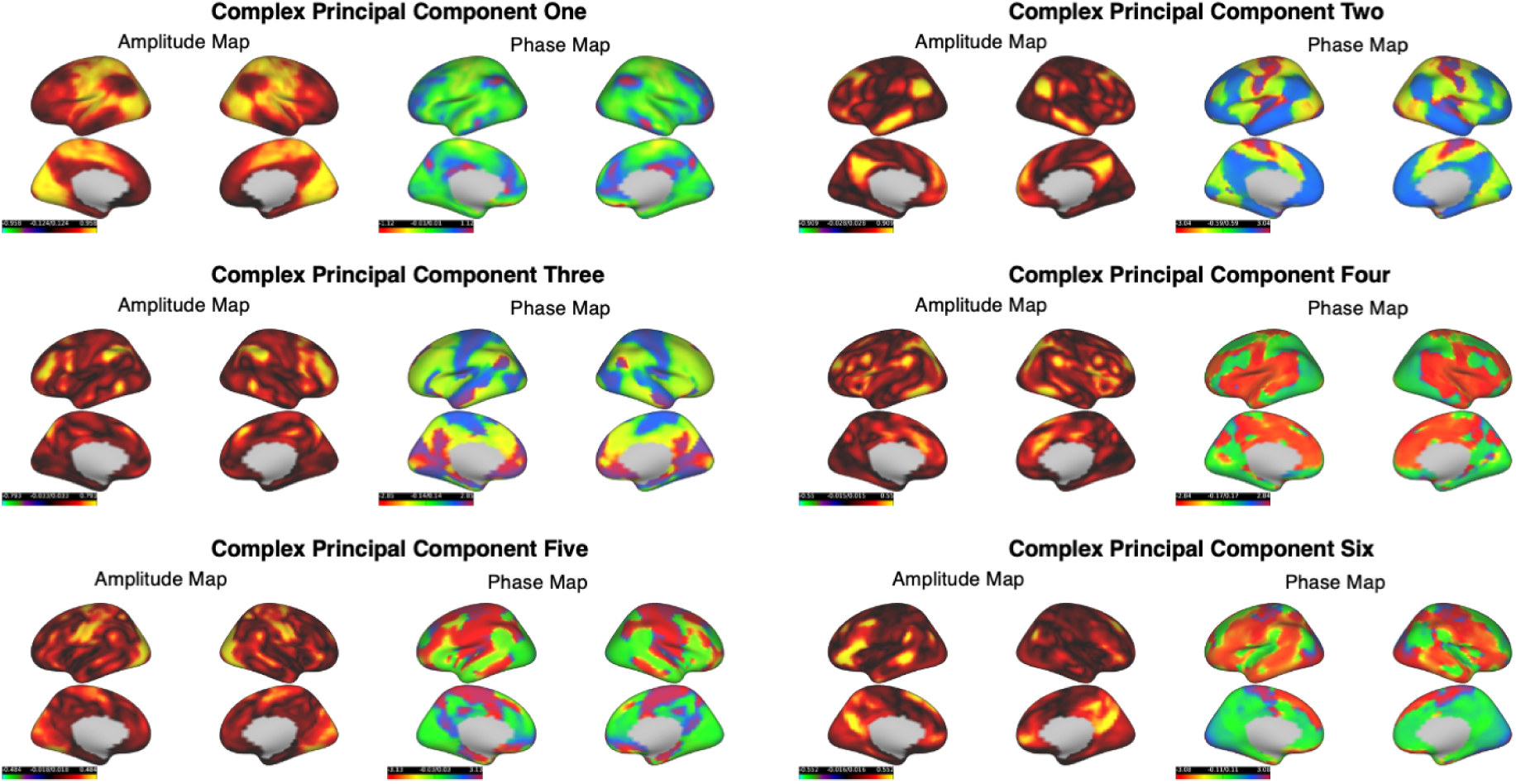
Complex Principal Components Beyond Three. **Amplitude and Phase Delay Maps of First Six Complex Principal Components.** While the application of the scree plot criterion identified three components as an optimal solution, there is still a large portion of variance left unexplained in BOLD time courses (∼68%). To further explore this remaining variance, we present the phase delay and amplitude maps of the first six complex principal components. Phase delay values are displayed in radians.

**Supplementary Figure 6.**
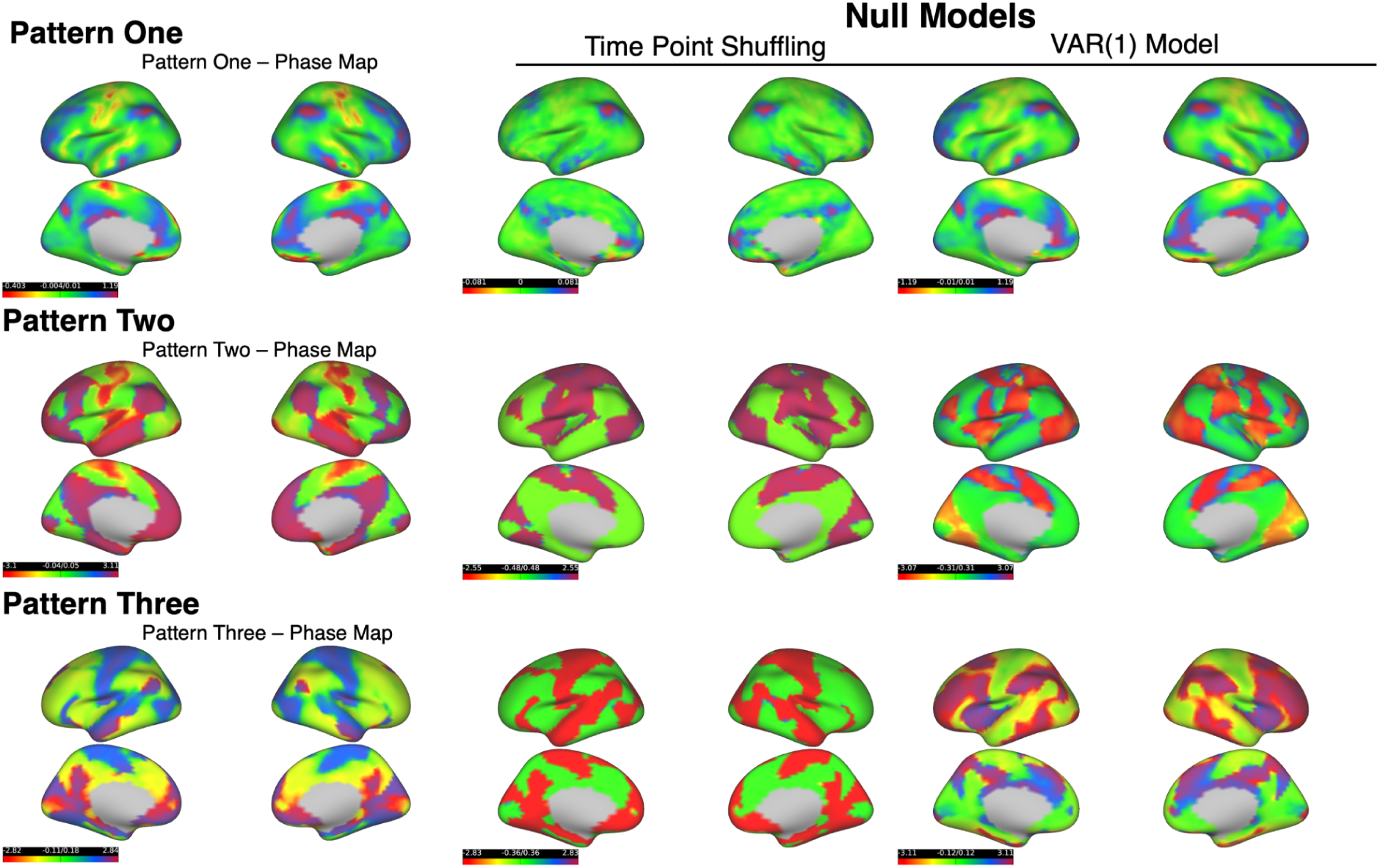
Complex PCA on Shuffled BOLD Time Courses. **Complex PCA Of Null Models.** One means to understand the properties of the cortical BOLD time courses that give rise to these spatiotemporal patterns is null modeling. By selectively preserving certain features of the time courses, we can identify the statistical properties of the time courses that are necessary for describing the spatiotemporal patterns. We conducted two null model exercises. First, to illustrate that the spatiotemporal patterns depend on properties of the time courses beyond zero-lag correlation, we can randomly shuffle the time points of the time courses; this procedure selectively preserves the zero-lag correlation structure of spontaneous BOLD fluctuations and removes the time-lag (autocorrelation) structure. We randomly shuffled the group-concatenated BOLD time courses and estimated three complex principal components from CPCA. We display the phase delay maps of the first three complex principal components from the shuffled time courses (middle) and original time courses (left) (displayed in radians). Random shuffling of BOLD time courses effectively eliminates the time-lag structure present in the original phase delay maps of the first three complex principal components. What is preserved is the zero-lag structure, i.e. in-phase and anti-phase statistical dependence. Visually, the phase delay maps of the shuffled BOLD time courses display a strictly bi-modal phase structure, corresponding to the anti-correlated fluctuations of the standing wave components of the complex principal components. The intermediate phase values present in the original complex principal components, corresponding to the traveling wave components, are eliminated. Second, we simulated time courses from a first-order multivariate autoregression model - i.e. VAR(1) - that was fit to the cortical BOLD time series. This model leaves the time-lag (i.e. autoregressive) structure of the cortical BOLD time courses intact, while assuming Gaussianity, linearity and stationarity of the time courses ^23^ . cPCA of the simulated time courses generated from the VAR(1) model replicated both the zero-lag and time-lag structure of the three spatiotemporal patterns (right). These findings suggest that three spatiotemporal patterns arise from properties of the BOLD time courses that are (mostly) Gaussian, linear and stationary.

**Supplementary Figure 7.**
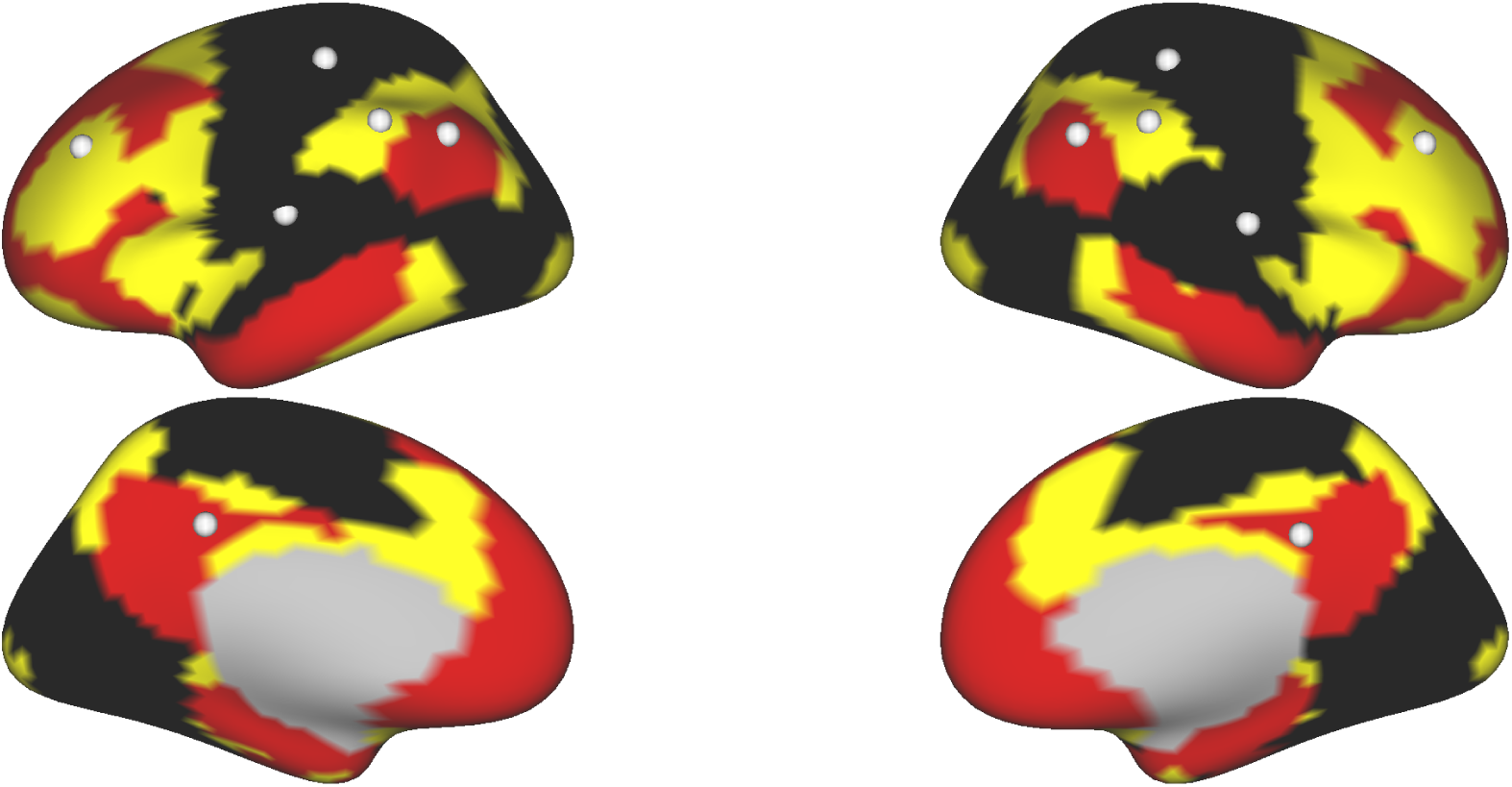
Location of Seed Regions in Seed-Based Regression and Co-Activation Pattern Analysis. **Location of Seed Regions in Seed-Based Regression and Co-Activation Pattern Analysis.** There are various methods for determining the location of seed regions. In our analysis, we chose seed regions within the three central networks of the three dominant spatiotemporal patterns - somatomotor (SM) and lateral visual (LV) networks, frontoparietal network (FPN) and DMN. The spatial outline of the SMLV, DMN and FPN for guiding the selection of seed regions were determined through a k-means clustering analysis of cortical vertices based on the similarity in their BOLD time courses. We found that a three-cluster k-means clustering solution precisely delineated the spatial outline of the three networks. Six bi-lateral seed locations are displayed for the SMLV (black) , FPN (yellow) and DMN (red). For the analyses in this study, we presented the results from seeds placed in the somatosensory cortex, precuneus and supramarginal gyrus. To test the robustness of our analyses to seed location, we also ran seed-based regression and CAP analyses with seeds placed medial insula (SMLV), inferior parietal cortex (DMN) and DLPFC (FPN). Because the results were found to be identical with the somatosensory cortex, precuneus and DLPFC, respectively, we do not present results for these seeds.

**Supplementary Figure 8.**
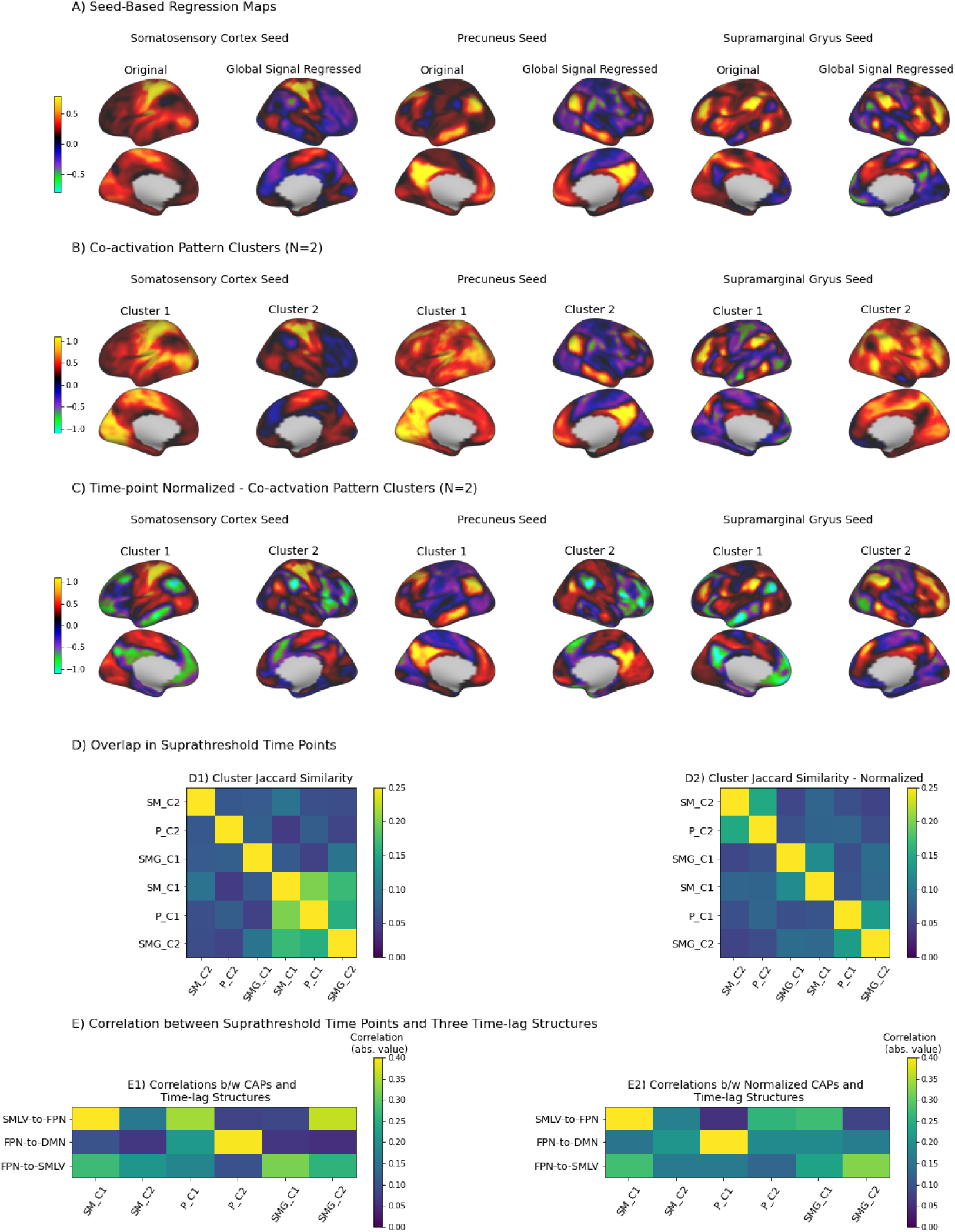
Seed-Based Analysis and Their Relationship to the Three Spatiotemporal Patterns. **Seed-Based Regression and CAP Analysis. (** SM = somatosensory cortex; P=Precuneus; SMG=Supramarginal Gyrus). FC topographies of seed-based regression maps and CAP centroids from somatosensory (SM), precuneus and supramarginal gyrus seeds. **A)** Seed-based regression maps with (left hemisphere) and without global signal regression (right hemisphere) for SM, precuneus and supramarginal gyrus seeds. We found that the effect of global signal regression is primarily a centering operation of correlation values, with the pattern of correlations largely consistent between the original and global-signal regressed correlation patterns - somatosensory (***r* = 0.96**), precuneus (***r* = 0.89**), supramarginal gyrus (***r* = 0.92**). For CAP analysis, we chose a threshold equal to the 85th percentile of the seed time course BOLD values, consistent with previous applications ^15^. **B)** CAP cluster centroids (N=2) from k-means clustering of non-normalized (i.e. not z-scored) suprathreshold time points from SM, precuneus and supramarginal seeds. **C)** CAP cluster centroids (N=2) of the same suprathreshold time points with normalization (i.e. z-scored) before input to the k-means clustering algorithm. Normalization (**Panel C**) results in two anti-correlated CAPs per seed, as opposed to a globally-positive vs. anti-correlated CAP per seed (**Panel B**) in the non-normalized solution. **D1)** Temporal overlap between binary time courses of the two CAPs from each seed using the Jaccard similarity (Jaccard index). The Jaccard similarity between two CAP binary time courses varies from 0 to 1, and reflects the ratio of overlapping onset time points (=1) to the total number of time points (N=60,000). To account for potential time-lags between CAP binary time courses, we took the maximum Jaccard similarity between the CAP binary time courses at a max temporal lag of 30 time points. Examination of the temporal overlap between CAP binary time courses revealed that the onsets of globally-positive CAP patterns (somatosensory C1, precuneus C1 and supramarginal C2) tended to co-occur at much greater rate than the anti-correlated CAPs (global CAPs: *=* 0.173 > anti-correlated CAPs: *=* 0.073). This is consistent with the global synchronization effect associated with the global mean time course. **D2)** Temporal overlap between CAP binary time courses from the normalized solutions of each seed analysis. The temporal overlap observed between globally-positive CAP time points in the non-normalized solution disappears in the normalized solution. **E)** Temporal correlation between the time course of the three spatiotemporal patterns (patterns one, two and three) and the CAP binary time courses for the non-normalized (**E1**) and normalized (**E2**) solutions. We found that CAPs with globally-positive BOLD activation patterns (**E1)** somatosensory C1, precuneus C1 and supramarginal C2) were most strongly correlated with the time course of pattern one: somatosensory cortex (SM; ***r* = 0.49**), precuneus (***r* = 0.35**), and supramarginal gyrus (***r* = 0.37**). The temporal correlation between the pattern one and normalized CAPs across the three seeds is reduced (**E2**), excluding the somatosensory CAP (C1).

**Supplementary Figure 9.**
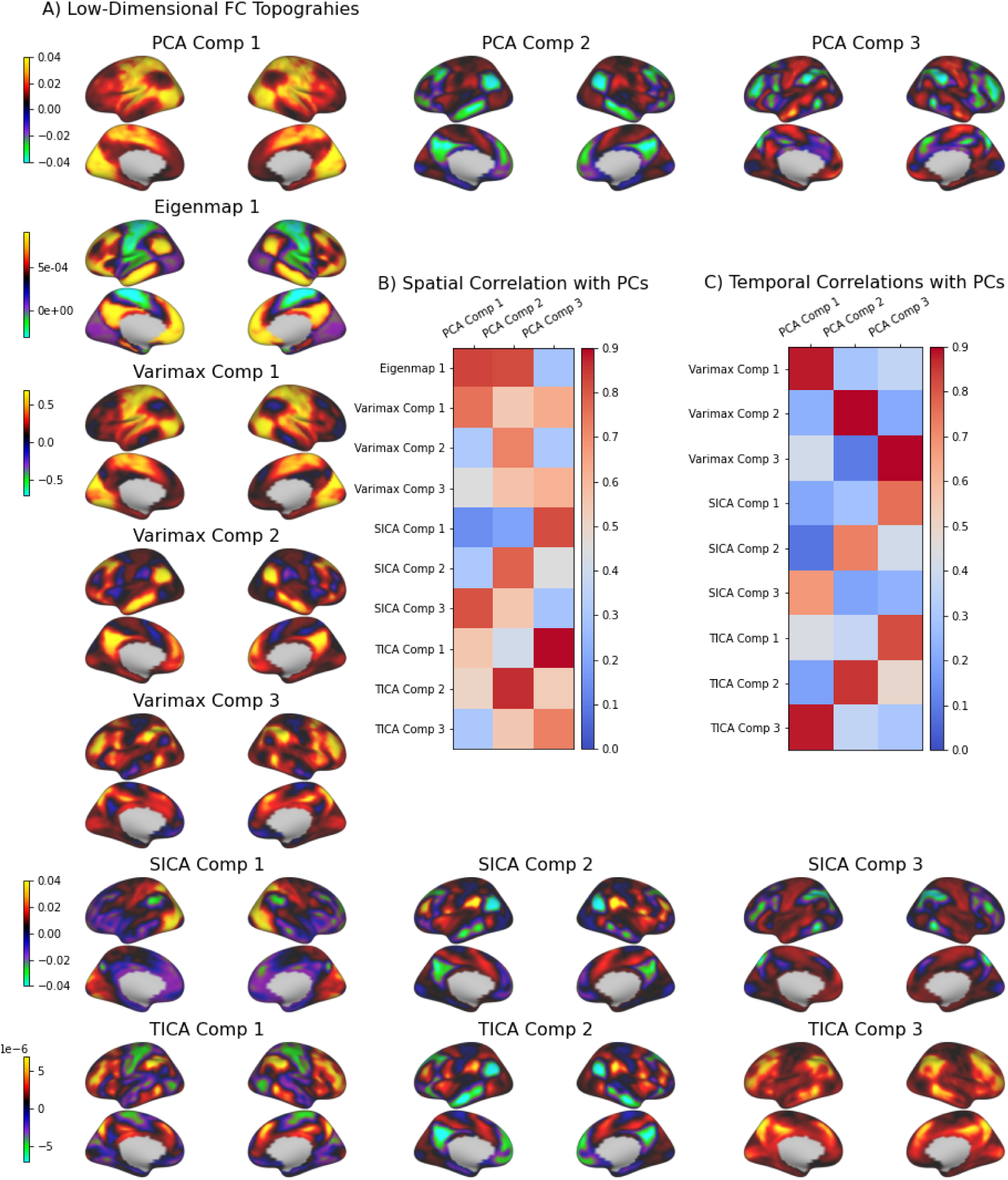
Dimension Reduction Functional Connectivity Topographies. **Functional Connectivity Topographies of Dimension Reduction Analyses.** (SICA=Spatial ICA; TICA = Temporal ICA). **A)** The spatial weights of components from PCA (N=3), Laplacian Eigenmaps (N=1), varimax rotation of principal components (N=3), spatial ICA (N=3) and temporal ICA (N=3). The temporal and spatial correlations (absolute value) between the components of dimension-reduction analyses and the first three principal components are shown in the middle of the plot. Note, due to the nature of the Laplacian Eigenmap algorithm as a non-linear manifold learning algorithm, time courses cannot be extracted for their components. In relation to principal components derived from PCA, spatial and temporal ICA amounts to a rotation (i.e. unmixing matrix) of the whitened temporal or spatial principal component axes such that the statistical independence between the axes is maximized, respectively ^31,72^. In other words, ICA rotates the original PCA solution to maximize a different criterion: statistical independence in the temporal or spatial domain. In this conception, ICA is one of a larger family of principal component rotation methods that also includes rotations towards so-called ’simple structure’. Simple structure rotations rotate the principal component loadings so that the parsimony of the loadings are maximized (each vertex loads strongly on only one component). We chose a popular simple structure rotation, Varimax rotation - an orthogonal rotation of the principal component loadings that maximizes simple structure. It is important to emphasize that the total cumulative variance explained by the principal component axes remains the same before and after rotation. As illustrated in the spatial and temporal correlations table, the dimension-reduction analyses are largely consistent in their spatial topographies and temporal dynamics with the first three principal components. In all cases, the rotation methods (SICA, TICA and Varimax) return components more or less similar in spatial and temporal dynamics to the first three principal components. In other words, despite the differing mathematical assumptions and objective criteria of these dimension-reduction methods, the results produced from each method for low-dimensional solutions are roughly consistent.

**Supplementary Figure 10.**
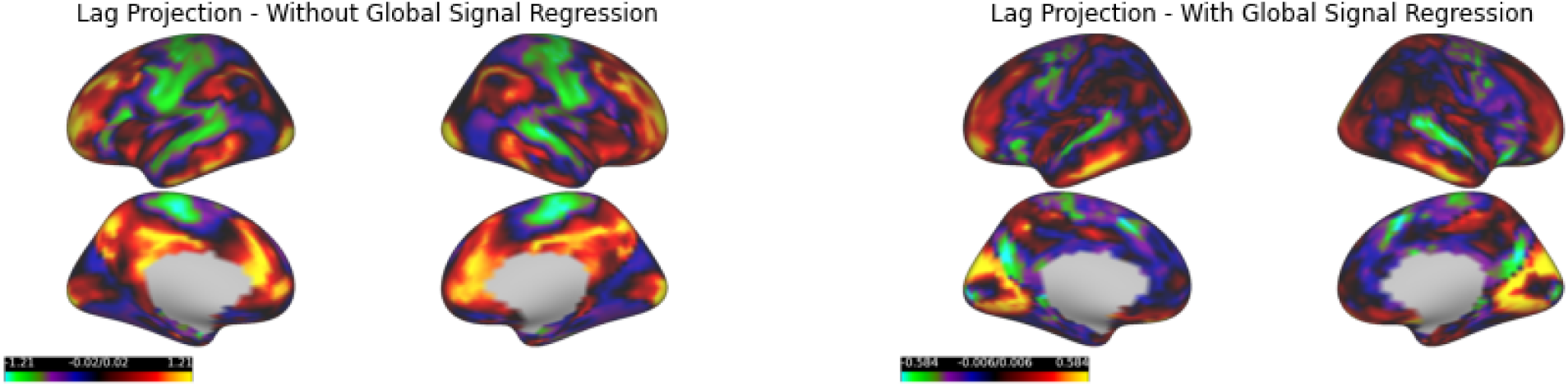
Global Signal Regression and Lag Projections. **Comparison of Lag Projections With and Without Global Signal Regression.** The lag projection from our study partially resembles the group average lag projection observed in Mitra et al. ^18^. However, our data differs in one important respect: Mitra et al.^18^ performed global signal regression as a preprocessing step. We display lag projections with and without global signal regression as a preprocessing step. Values on each cortical map represent the average time-delay between each cortical vertex and all others. Time-delay values are colored from light green/blue (earlier in time) to bright yellow/green (later in time). The range between the earliest and latest time-delay values are significantly shorter for lag projections on global-signal regressed data. The lag projection of the global signal regressed data resembles the spatial distribution time-lags observed in Mitra et al.^18^: BOLD activity beginning in superior medial prefrontal cortex, inferior precuneus, motor cortex, anterior cingulate cortex, and temporal gyrus and ending in the DMN and visual cortex. In addition, the length of the lag projection is now 1 sec (cut in time by half from non-globally regressed data), closely matching the duration found by Mitra et al. This is consistent with the observation by Mitra et al. that global signal regression reduces the range of observed latencies between BOLD time courses.

**Supplementary Figure 11.**
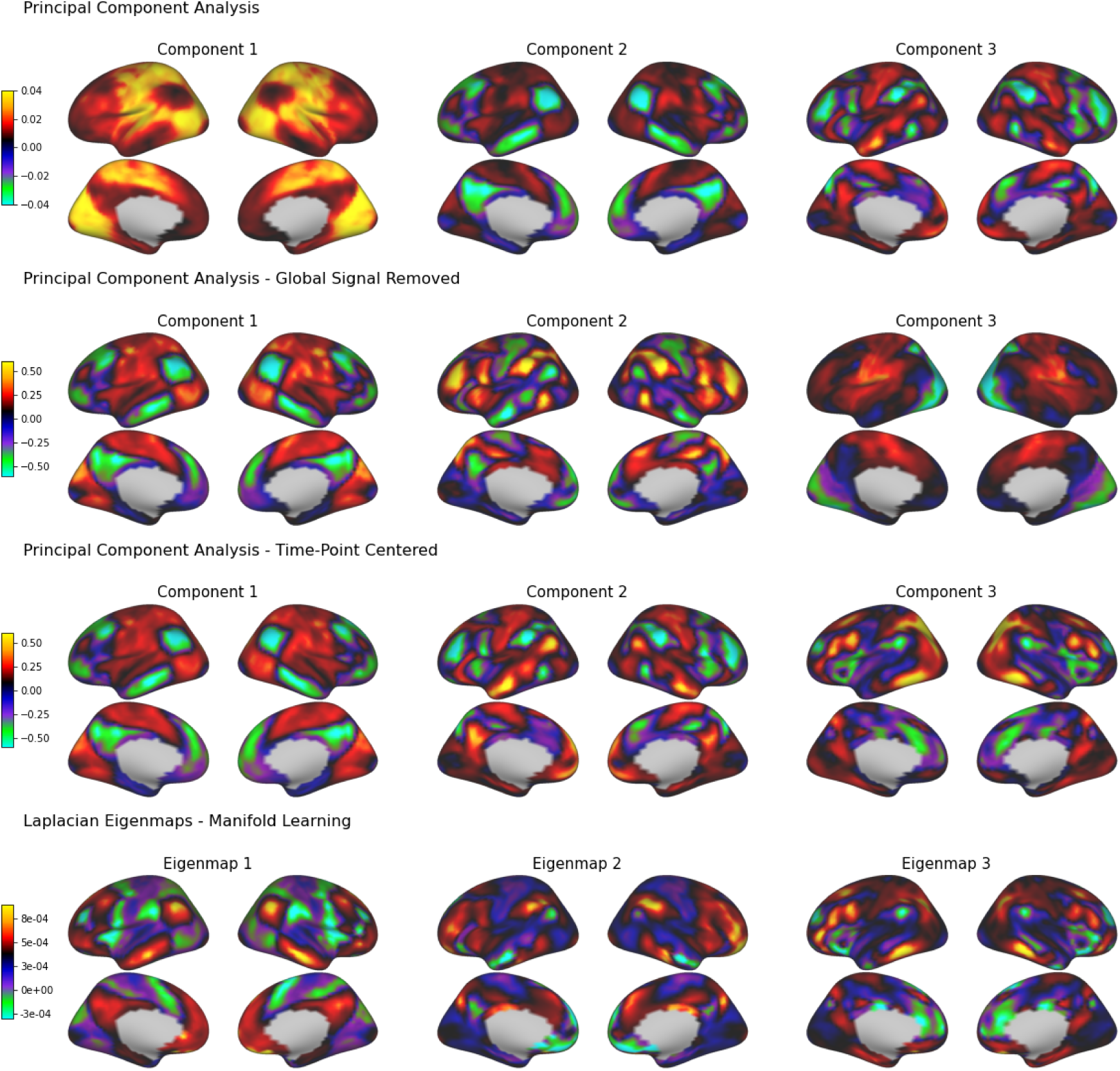
PCA and Functional Connectivity Gradients. **Principal Component and Laplacian Eigenmap Topographies.** Displayed are the FC topography spatial weights from PCA, PCA on global-signal regressed data, PCA on time-point centered data, and Laplacian Eigenmaps (LE). Note, LE analysis with a radial basis function kernel (non-linear kernel) was also tried and the results were very similar. Note, we observed that the eigenmaps were highly positively skewed. To make the negative values of the eigenmaps more visible the colormap is made non-symmetric.The first eigenmap corresponds to the principal functional connectivity gradient (PG)^2^. However, we note that the exact spatial pattern of the PG depends on the level of thresholding applied to the FC matrix (Figure 5). In this LE solution, no thresholding was applied. The first and second Laplacian Eigenmaps match the second and third principal components from the PCA solution, respectively (with an arbitrary sign-flip). The difference between the PCA and LE solution is that the first principal component seems to be missing from the LE solution. However, the first three components from the PCA on global-signal regressed data and time-point centered do match the three Laplacian Eigenmaps (with an arbitrary sign flip). These similarities between the spatial maps produced by PCA and Laplacian Eigenmaps have been previously observed by Vos de Waal et al.^25^. An important question is why the first principal component, the variance upon which BOLD time courses vary the greatest, is not returned by LE and PCA applied to global-signal regressed and time-point centered data? In all three cases, the difference is due to the same mechanism: mean-centering along the time domain (i.e. mean centering vertex BOLD values within a time point). Pattern one is precisely tracked by the global signal (***r* = 0.97**). Time-point centering and global signal regression have similar effects - reducing or eliminating the variance of the global signal time courses^15^. In the Laplacian Eigenmap solution, a time-point centering is not as explicit. Manifold learning of a time vertex-by-vertex kernel matrix operates on a mean centering of the feature space, i.e. each time point is centered. As illustrated, this has the same practical effect as global signal regression and time-point centering. Thus, LE, PCA of global-signal regressed data and PCA of time-point centered data return very similar spatial patterns (some with an arbitrary sign difference).

**Supplementary Figure 12.**
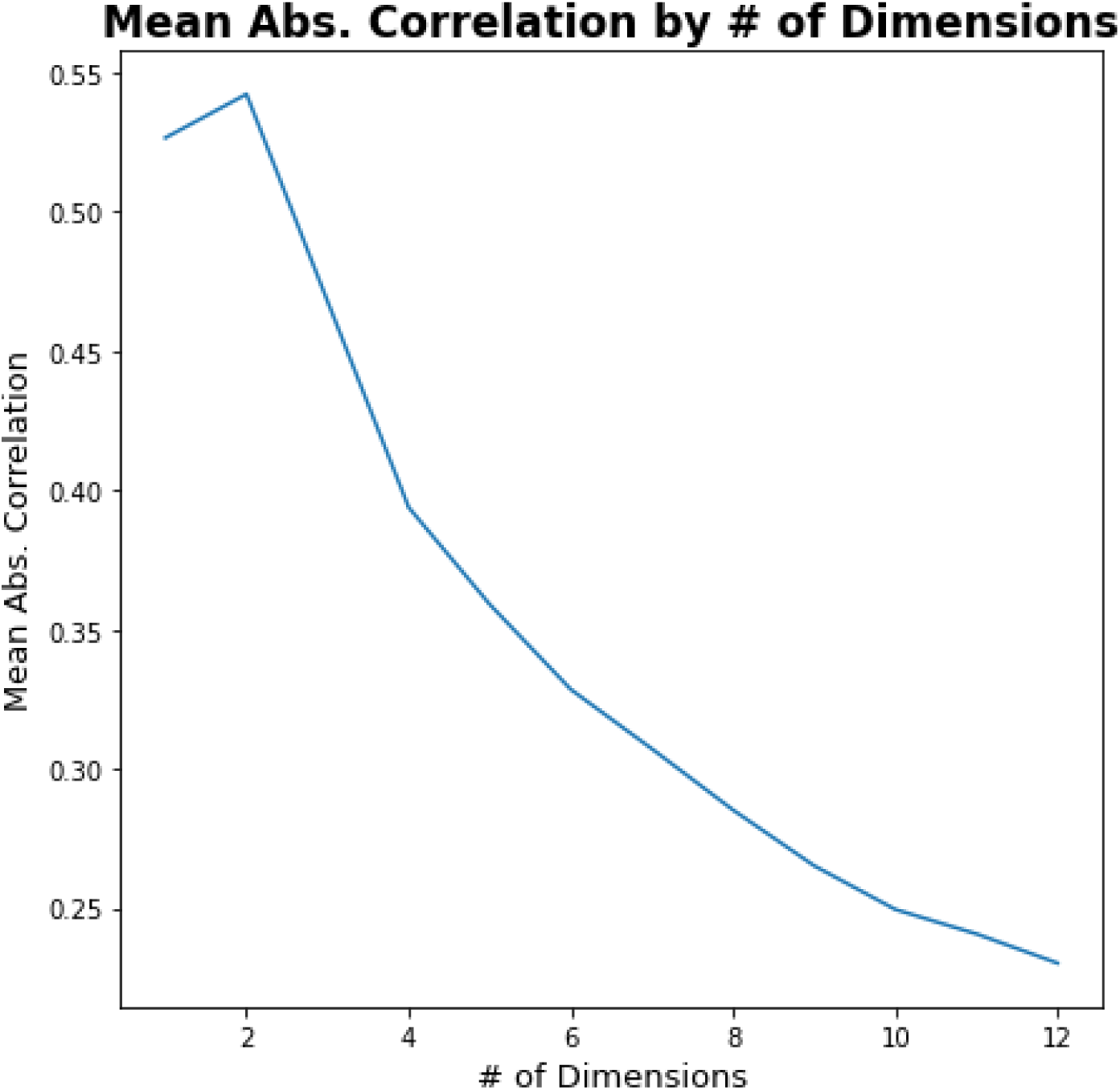
Consistency in Zero-lag FC Topographies at Finer-Grained Solutions. **Consistency in FC Topographies by Granularity of Solutions.** One of the findings of our study is that there is a surprising consistency in FC topographies estimated from ‘three-dimensional’ zero-lag analyses. However, it is worth exploring the degree of consistency in FC topographies at higher component numbers. We estimated higher number component (cluster) solutions for several latent dimension-reduction and clustering methods - including PCA, varimax-rotated PCA, temporal ICA, spatial ICA, HMM, and CAP analysis (all three seeds from main text). We examined the consistency in FC topographies at component numbers varying from one to 12. Note, the same number of components (clusters) was estimated across methods at each value. While it is difficult to quantify the degree of consistency in outputs from multiple methods with a single number, we used the mean absolute-valued correlation between all pairs of FC topographies produced by all methods as a rough estimate. We display the mean absolute-valued correlation by component number solution. As illustrated in the plot, the consistency of FC topographies drops significantly at higher component (cluster) numbers.

**Supplementary Figure 13.**
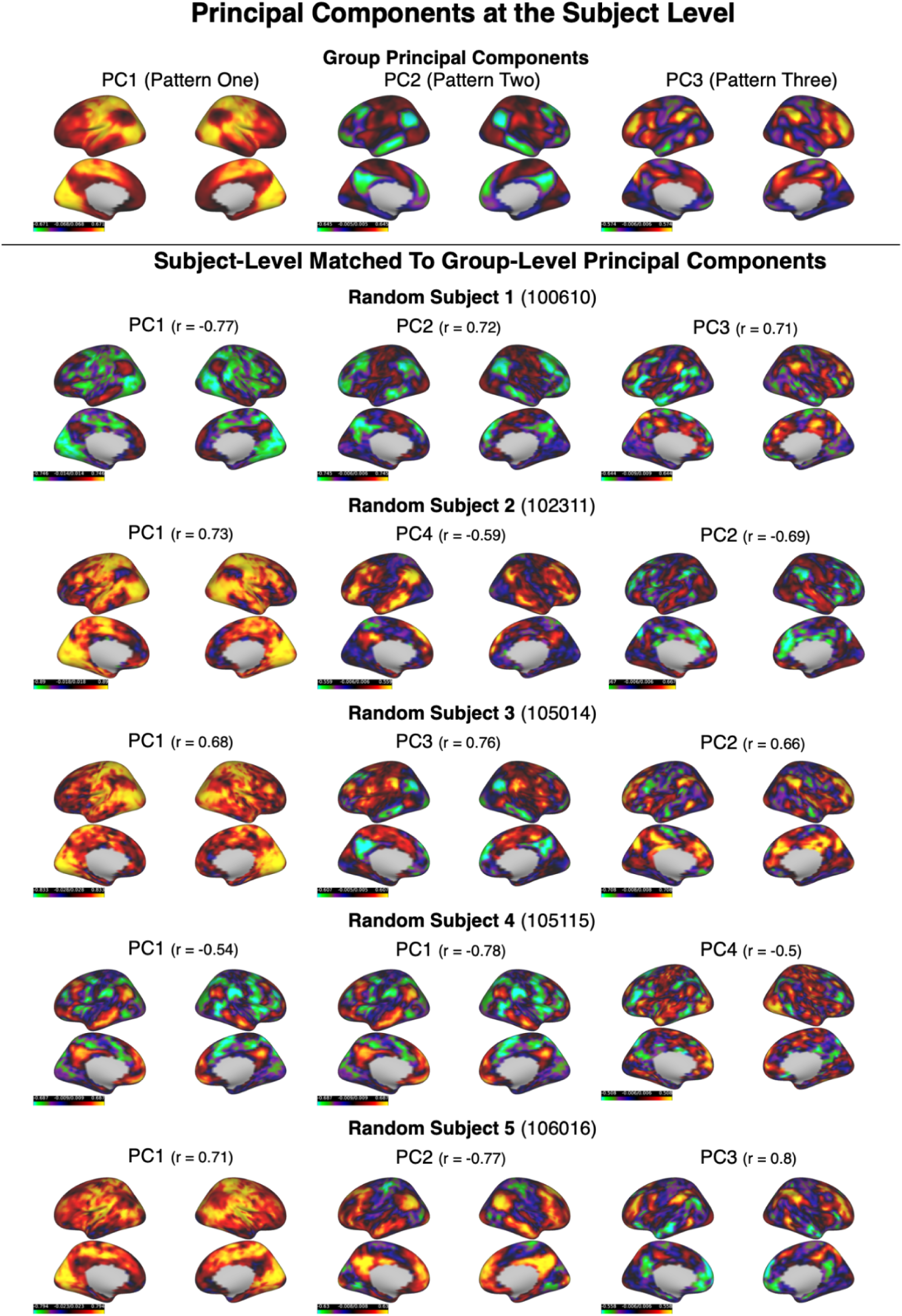
Inter-Subject Variability in Principal Components. **Intersubject Variability of Spatiotemporal Patterns.** To assess the degree of intersubject variability in the three spatiotemporal patterns we randomly selected five subjects from our group analysis for further study. As illustrated in Figure 3, PCA effectively captures the zero-lag correlation structure of the three spatiotemporal patterns (patterns one, two and three). We conducted PCA on the BOLD time courses of the five subject scans separately, and compared the spatial structure of the first five principal components to the first three group-level principal components. As the ordering of the subject-level principal components by explained variance may not correspond precisely to the ordering of the group-level principal components, we conducted a matching procedure to map the subject-level principal components to the first three group-level principal components. This matching procedure was guided both by the spatial correlation between the subject-level and group-level principal components and by visual comparison. Beside each subject’s header is their corresponding HCP ID. For each subject, their matching principal component was displayed in the same column as the group-level principal component (PC1: left, PC2: middle, PC3: right). Beside the header of each subject’s principal component is the spatial correlation between that principal component and its matched group-level principal component (in the same column). Overall, patterns one, two and three were present in all five subjects, with notable intersubject variability. Several types of variability were observed: 1) the ordering by explained variance of the subject-level principal components was sometimes different than that observed at the group-level, 2) the same spatiotemporal patterns exhibited global and anti-correlated types across subjects, and 3) two subject-level principal component matched most closely to a single spatiotemporal pattern (for one subject - subject 4).

## Methods and Materials

### Resting-State fMRI Data

Our study utilized resting-state fMRI scans from the Human Connectome Project (HCP) S1200 release^47^. Participants were unrelated, healthy young adults (ages 22–37). Resting-state fMRI data was collected over two consecutive days for each subject and two sessions, each consisting of two 15 minute runs, amounting to four resting-state scans per subject. Within a session, the two runs were acquired with opposite phase encoding directions: L/R encoding and R/L encoding. We selected a single 15 min scan from a random sample of participants (n=50; 21 males) on the first day of scanning. We balanced the number of L/R and R/L phase encoding scans across our participants (n=25 for each encoding direction) to ensure results were not biased by acquisition from any given phase encoding direction. We chose a single 15 min scan per participant to ensure that the phase encoding/decoding parameter and the imaging session (two resting-state scans per imaging session) did not differ within the same participant. A second independent random sample of participants (n=50, 22 males) was used as a validation sample. We selected surface-based CIFTI resting-state fMRI scans (MSMall registered) that had been previously preprocessed with the HCP’s ICA-based artifact removal process^48^ to minimize effects of spatially structured noise in our analysis. All brain-imaging data were acquired on a customized Siemens 3 T Skyra at Washington University in St. Louis using a multi-band sequence. The structural images were 0.7 mm isotropic. The resting-state fMRI data were at 2 mm isotropic spatial resolution and with TR = 0.72 s temporal resolution. Further details of the data collection and preprocessing pipelines of the HCP can be found elsewhere ^47,48^. Informed consent was obtained from all subjects. All methods were carried out in accordance with relevant guidelines.

### Resting-State fMRI Preprocessing

Resting-state fMRI scans were spatially smoothed with a 5mm FWHM kernel using the surface-based smoothing algorithm in Connectome Workbench Version 1.4.2. Resting-state fMRI signals from each vertex were then temporally filtered to the conventional low-frequency range of resting-state fMRI studies using a Butterworth bandpass zero-phase filter (0.01-0.1Hz). Due to 1) the computational complexity of our analytic pipeline, owing to the large number of analyses studied, and 2) our interest in global, spatially distributed patterns, resting-state fMRI scans were then resampled to the *fs4* average space from Freesurfer^49^. This step down-sampled the total number of vertices in the left and right cortex to 4800 vertices. In group analyses, we z-scored (to zero mean and unit variance) the BOLD time series from all vertices before temporal concatenation of individual scans. All analyses were applied to group-level data formed by temporal concatenation of subject resting-state scans.

### Complex Principal Component Analysis

To extract traveling wave patterns, we apply PCA to complex BOLD signals obtained by the Hilbert transform of the original BOLD signals. We refer to this analysis as complex PCA (CPCA). This technique has been referred to as complex Hilbert empirical orthogonal functions in the Atmospheric and Climate sciences literature^50^, or complex orthogonal decomposition in the engineering/physics literature^14^.

CPCA allows the representation of time-lag relationships between BOLD signals through the introduction of complex correlations between the Hilbert transformed BOLD signals. The original time courses and their Hilbert transforms are complex vectors with real and imaginary components, corresponding to the non-zero-lagged time course (t=0) and the time course phase shifted by t=pi/2 radians (i.e. 90 degrees), respectively. The correlation between two complex signals is itself a complex number (composed of a real and imaginary part), and allows one to derive the phase offset (and magnitude) between the original time courses - i.e. the time-lag at which the correlation is maximum. CPCA applied to the complex-valued correlation matrix produces complex spatial weights for each principal component that can give information regarding the time-lags between BOLD time courses. In the same manner that a complex signal can be represented in terms of amplitude and phase components (via Euler’s transform), the real and imaginary components of the complex principal component can be represented in terms of amplitude and phase spatial weights. Of interest in this study is the phase delay spatial map that represents the time-lag between pairs of BOLD time courses - i.e. those cortical vertices with a low phase value activate earlier than cortical vertices with a high phase value. Importantly, the principal components from the CPCA retain the same interpretive relevance as the original PCA - the first N principal components represent the top N dimensions of variance in the Hilbert transformed BOLD signals. CPCA was implemented with singular value decomposition of the groupwise temporally-concatenated complex-valued time series using the fast randomized SVD algorithm developed by *Facebook* (https://github.com/facebookarchive/fbpca).

### Estimating The Time-Scale of Complex Principal Components

For simplicity, the phase spatial maps of each complex principal component are displayed in seconds (**Figure 2**), as opposed to radians. However, the conversion of phase values (in radians) to time-units (seconds) requires an estimation of the time-scale of each complex principal component. The phase spatial maps of the complex principal components have no characteristic time scale other than that imposed by our band-pass filtering operation (0.01 - 0.1 Hz, i.e. 100 to 10s) in the preprocessing stage. To approximate a unique time scale within this frequency range for each component, we calculated the average duration for a full oscillation of each complex principal component using the temporal phase of the complex component time series. This was calculated by fitting a linear curve to the unwrapped temporal phases of the complex principal component time series. The slope of the curve was then used as an estimate of the average duration in radians of a TR (0.72s) or time-points. To estimate the average duration in TRs of a full oscillation, we divided a full oscillation (2 radians) by the duration in radians of a TR. For example, for a TR duration of 0.5 radians, the duration of a full oscillation (2pi radians) would be approximately 12.6 TRs. Using this procedure, we found that the average duration of the first three complex principal components are ∼28s (38.7 TRs), ∼27s (37.4 TRs) and ∼28s (39.1 TRs), respectively. Using this duration as an estimate of the characteristic time scale of each complex principal component, allows us to provide an estimate of the time-delay in seconds of the spatial phase map. For example, for the first complex principal component, a 360 degree (2pi radians) phase difference between two cortical BOLD time series would correspond to a ∼28s time-lag between the time series. A smaller phase difference between two cortical BOLD time series, such as 1 radian, would correspond to ∼14s time-lag between the time series, and so forth.

### Temporal Reconstruction of Complex Principal Components

To examine the temporal progression of each complex principal component, we sampled the reconstructed BOLD time courses from each complex principal component at multiple, equally-spaced phases of its cycle (N=30; **Figure 2**). For each complex principal component, the reconstruction procedure was as follows: 1) the complex principal component time series was projected back into the original vertex-by-time space to produce time courses of the complex principal component at each vertex, 2) the temporal phase of the complex principal component time course was segmented into equal-width phase bins (N=30) spanning a full oscillation of the spatiotemporal pattern (0 to 2pi radians), and 3) the vertex values within each bin were averaged to produce a ‘snapshot’ of BOLD activity at each phase bin (N=30) of the spatiotemporal pattern. The end result is a spatiotemporal representation of each complex principal component in terms of time-varying BOLD activity at equally spaced phases of its cycle.

### Traveling Index of Complex Principal Components

The real and imaginary parts of a complex principal component correspond to the spatial weights of the component at zero and pi/2 (90 degree) phase shift of the original time courses. In a sense, they encode the temporal evolution of the complex principal component from one configuration (the real part) to a subsequent configuration (the imaginary part). By definition, a pure standing wave would exhibit the same spatial configuration from zero to pi/2 (90 degree) phase shifts of its cycle. A pure traveling wave would exhibit a different spatial configuration from zero to pi/2 (90 degree) phase shifts of its cycle. This observation suggests a means to quantify the degree of ‘traveling’ wave behavior of a complex principal component using the statistical dependence between its real and imaginary parts. A coherent traveling wave (i.e. propagation) of BOLD amplitudes across the cortex would exhibit one spatial configuration at one point in time and a different spatial configuration at another point in time. Thus, a complex principal component that encodes this traveling wave behavior would exhibit differing spatial configurations in its real and imaginary spatial weights. Utilizing a metric developed by Feeny ^14^, we define the ‘traveling’ index of a complex principal component as the reciprocal of the condition number of the matrix whose two columns are the real and imaginary parts of the complex principal component. This metric simply encodes the statistical dependence between the real and imaginary parts of the complex principal component. Pure traveling waves would exhibit completely orthogonal real and imaginary parts, and a traveling index of one. Pure standing waves would exhibit completely dependent real and imaginary parts, and a traveling index of zero.

### Analysis of Null Models

We performed two null model exercises to determine the statistical properties of the cortical BOLD time courses that are necessary to capture the time-lag structure of the complex principal components. Null models allow the selective removal of features of the time courses, such as removing autocorrelation structure and preserving zero-lag correlation structure. First, to illustrate that the spatiotemporal patterns depend on properties of the time courses beyond zero-lag correlation, we randomly shuffled the time points of the time courses; this procedure selectively preserves the zero-lag correlation structure of spontaneous BOLD fluctuations and removes the time-lag (autocorrelation) structure. We randomly shuffled the group-concatenated BOLD time courses and estimated three complex principal components from cPCA. Second, we simulated time courses from a first-order multivariate autoregression model - i.e. VAR(1) - that was fit to the cortical BOLD time series. A VAR(1) model predicts each time course from preceding time points (lag of one time point) of itself and all other time courses (i.e. all cortical vertices - 4800 time courses). This model leaves the time-lag (i.e. autoregressive) structure of the cortical BOLD time courses intact, while assuming Gaussianity, linearity and stationarity of the time courses ^23^ . Due to the large number (4800 cortical vertices) of highly collinear time courses, we extracted 200 dimension-reduced time courses from the original time courses using PCA. The VAR(1) model was fitted on the PC time courses, and simulated time courses (the same length as the original time courses) were generated from the fitted model. The simulated PC time courses were then projected back into the original cortical vertex space. CPCA was then applied to these simulated time courses.

### Zero-Lag Functional Connectivity Analyses

#### Description of Zero-lag FC Analyses

Following the standard terminology of the functional magnetic resonance imaging (fMRI) literature, we refer to zero-lag synchrony among intrinsic BOLD fluctuations as ‘functional connectivity’ (FC) ^51^. FC between cortical brain regions organize into global, cortex-wide patterns, referred to as ‘FC topographies’. All analyses were conducted so as to be consistent as possible with previous studies. For some of these analyses, results were compared with and without global signal regression. Global signal regression was performed by regression of the global mean time series (averaged across all cortical vertices) on all cortical time series. Residual time series from each regression were then used for subsequent analysis. All analyses were conducted using custom Python scripts, and are publicly-available at https://github.com/tsb46/BOLD_WAVES. The following zero-lag FC analyses were conducted:

- Principal component analysis (PCA): consists of eigendecomposition of the empirical covariance matrix of the vertices’ time series, or alternatively, singular value decomposition of the mean-centered group data matrix (time series along rows, vertices as columns). The first *T* principal components represent the top *T* dimensions of variance among cortical BOLD time courses. By construction, the first principal component is the latent direction of variation with the largest explained variance across all input variables, followed by the second most explanatory component, and so forth. The principal component spatial weights on each vertex were used to interpret the spatial patterns of each principal component. Principal component scores were obtained from the projection of the temporally-concatenated group time series onto the principal component space, and represent the time course of each principal component.
- Varimax rotation of principal components: consists of an orthogonal rotation of the principal component spatial weights, such that the simple structure of the spatial weights are maximized. Simple structure is defined such that each vertex loads most strongly one component, and weakly on all others. We adapted code from the Python package *xmca* (https://github.com/nicrie/xmca) for implementation of varimax rotation.
- Laplacian Eigenmaps (spectral embedding): is a nonlinear manifold learning algorithm popular in the FC gradient literature^25^. The input to the Laplacian eigenmaps algorithm was the vertex-by-vertex cosine similarity matrix^2^, representing the similarity in the BOLD time series between all cortical vertices. Of note, cosine similarity is equivalent to Pearson correlation in mean-centered and unit normalized time series (i.e. z-score normalization), as was the case with our data. Laplacian Eigenmaps performs an eigendecomposition of the transformed similarity matrix, known as the normalized Laplacian matrix. We also computed Laplacian Eigenmaps with a Gaussian radial basis function (gamma=1), and the results were virtually identical to the cosine similarity metric. We used the spectral embedding algorithm implemented in the *scikit-learn* Python package, and details can be found at (https://scikit-learn.org/stable/modules/generated/sklearn.manifold.SpectralEmbedding.html).
- Spatial and temporal independent component analysis (ICA): estimates linearly mixed, statistically independent sources from a set of input variables. In the case of spatial ICA, principal component axes derived from PCA of the *time point-by-time point covariance matrix* are rotated to enforce statistical independence in the spatial domain. In the case of temporal ICA, principal component axes derived *from PCA of the vertex-by-vertex covariance* are rotated to enforce statistical independence in the temporal domain. As with varimax rotation, we input a three principal component solution for both temporal and spatial ICA. We used the FastICA algorithm implemented in the *scikit-learn* Python package. Details can be found at (https://scikit-learn.org/stable/modules/generated/sklearn.decomposition.FastICA.html).
- Seed-based correlation analysis: consists of correlations between a seed brain region time course and time courses of all cortical vertices. Seed-based correlation analysis was performed for three seed locations. There are various methods for determining the location of seed regions. In our analysis, we chose seed regions within the three most prominent networks in the three prominent spatiotemporal patterns - SMLV, FPN and DMN. We chose seeds in the somatosensory cortex (SMLV), precuneus (DMN), and supramarginal gyrus (FPN) (**Supplementary Figure 7)**. The spatial outline of the SMLV, DMN and FPN for guiding the selection of seed regions were determined through a k-means clustering analysis of the temporally-concatenated group time series with cortical vertices as observations and BOLD values at each time points as input variables (i.e. features). We found that a three-cluster k-means clustering solution precisely delineated the spatial outline of the three networks. This spatial outline was used to ensure the seeds were placed within their appropriate location of each network. In addition, we also tested the robustness of our results for different seed locations in the three networks - medial insula (SMLV), inferior parietal cortex (DMN) and dorsolateral prefrontal cortex (FPN) - and found that the results were identical.
- Co-activation pattern (CAP) analysis: Three CAP analyses were performed for the same three seed regions used in the seed-based regression analysis. CAP analyses first identify time points with the highest activation for a seed time course. Consistent with previous studies^15^, we chose the top 15% of time points from the seed time course. The BOLD values for all cortical vertices in the top 15% time points are then input to a k-means clustering algorithm to identify recurring CAPs of BOLD activity. We chose a two cluster solution for all CAP analyses. For each seed, the two cluster centroids from the k-means clustering analysis represent two CAPs associated with the seed time course.
- Hidden Markov modeling (HMM): is a probabilistic generative model used to infer the sequence and form of discrete hidden states, as well as their transition probabilities from an unobserved sequence of latent states. HMM construes the data-generating process based on multivariate Gaussian distributions conditioned on unknown latent ‘brain states’ that are assumed to generate the observed cortical BOLD time series. Each brain state represents a recurring pattern of BOLD co-activations/deactivations, somewhat similar to CAPs. To avoid overfitting and to reduce noise in the high-dimensional input data, we conducted a PCA of the cortical BOLD time series. The first 100 principal component projections of the time series served as input to the HMM algorithm. Associated with each brain state is a mean amplitude vector with a value for each principal component (N = 100), and a covariance matrix between the 100 principal component time courses. The mean amplitude vector represents the pattern of BOLD activity amplitudes associated with that brain state. For interpretation, the mean amplitude vector is projected back into cortical verex space for interpretation. A variety of potential ‘observation models’ are frequently used in HMM models. As cortical time series are measured on a continuous scale (as opposed to discrete measurements), the probability of a time point conditional on a hidden brain state (i.e. emission probabilities) is modeled as a mixture of Gaussian distributions. We used the HMM algorithm with Gaussian mixture emission probabilities implemented in the Python package *hmmlearn* (https://github.com/hmmlearn/hmmlearn).
- Modularity analysis: to determine whether the community structure of cortical BOLD time courses can be explained by the three spatiotemporal patterns, we applied the Louvain modularity-maximization algorithm to the original FC matrix and a FC matrix reconstructed from the three spatiotemporal patterns. FC matrices were calculated by computing the Pearson correlation coefficient between all pairs of cortical vertex time courses. The reconstructed FC matrix was created from projecting the timecourses of the three complex principal components back on to vertex space and computing a FC matrix. The Louvain modularity-maximization algorithm was applied with a resolution parameter value of 1 (with asymmetric treatment of negative weights). To compare the degree of similarity between the community structure of the original and reconstructed FC matrix, we computed the normalized mutual information (NMI) between their community assignments from the Louvain algorithm. The NMI varies from 0 (completely independent communities) to 1 (completely identical communities). This analysis was performed using the Python package *bctpy* (https://github.com/aestrivex/bctpy).

### Model Selection: Choice of Number of Dimensions in Dimension-Reduction Algorithms

The dimension-reduction algorithms used in this study, including PCA, PCA with varimax rotation, spatial and temporal ICA, and Laplacian Eigenmaps, as well as HMMs, require a choice of the number of latent dimensions/hidden states to estimate. For PCA with varimax rotation, spatial and temporal ICA, and HMM, this controls the degree of richness and/or fine-grained distinctions of the data description - i.e. how many separate unobserved hidden phenomena are assumed and quantitatively modeled to underlie each given data point or observation. We did not assume or try to derive a single ‘best’ number of latent dimensions to represent intrinsic functional brain organization ^52,53^. As we were interested in large-scale cortical patterns of FC, our survey focuses on low-dimensional latent solutions. As an initial estimate of the number of latent dimensions for all choices of dimension reduction algorithms, we examined the first *T* dominant axes of variation (i.e. principal components) of the correlation matrix formed between all pairs of cortical BOLD time series. Specifically, we examined the flattening (or diminishing return) in explained variance (i.e. eigenvalues) associated with neighboring principal components, a procedure known as Catell’s scree plot test^20^. According to this test, the number of components to extract is indicated by an ‘elbow’ in the plot, representing a ‘diminishing return’ in extracting more components. Clear elbows in the scree plot were observed after a principal component solution of one and three (**Figure 3C**). We chose the higher-dimensional solution of three components. Note, the elbow in explained variance after three components was independent of the functional resolution (i.e. vertex size) of the cortex - we found the same elbow after three components in a scree plot constructed from high-resolution functional scans (∼60,000 vertices without downsampling to 4,800 vertices as described above in our preprocessing pipeline). Thus, three latent dimensions were estimated for all dimension-reduction algorithms, and three hidden states were estimated for the HMM.

### Quasiperiodic Pattern and Lag Projections

There are two widely-used algorithms for the study of spatiotemporal patterns in BOLD signals: 1) interpolated cross-covariance functions (Mitra et al. ^7,54^) for the detection of lag/latency projections (∼0-2s) and 2) a repeated-template-averaging algorithm of similar spatiotemporal segments^18^ for detection of the QPP (∼20s).

Lag projections represent the average time-lag between a brain region’s time course and all other brain regions. It provides an estimate of the average temporal ‘ordering’ of brain region time courses, such that a brain region with a greater average time-lag occurs after a brain region with a smaller average time-lag. For our study, we applied the lag projection algorithm to all cortical vertex time courses. The time-lag between a pair of cortical vertex time courses is defined as the peak of their lagged cross-covariance function. Lag projections are derived as the column average of the pairwise time-lag matrix between all cortical vertex time courses. An analogous column-wise averaging operation can be applied to the complex correlation matrix used by CPCA. Specifically, we computed the column-wise circular mean of the pairwise phase-delay values of the complex correlation matrix. The circular mean was computed due to the circular nature of the phase values of a complex correlation (i.e. -pi and pi are identical phase differences).

To estimate the QPP, the template-autoregressive matching algorithm of Majeed et al. ^18^ was used. The algorithm operates in the following manner: start with a random window of BOLD TRs, compute a sliding window correlation of the window across the temporally concatenated group data at each time point, and then average this segment with similar segments of BOLD TRs (defined using a correlation threshold). This process is repeated iteratively until a level of convergence is reached. The result is a spatiotemporal averaged template of BOLD dynamics (that could be displayed in a movie, for example), along with the final sliding window correlation time series. The final sliding window time series is the same length as the original subject or group concatenated time series and provides a time index of the appearance of the QPP in BOLD data. Python code for this analysis was modified from the C-PAC toolbox (https://fcp-indi.github.io/). Consistent with previous studies ^18,28^, the following parameters were chosen for the template matching algorithm: the window length was 30 TRs, the maximum correlation threshold for identifying similar segments was *r >* 0.2, and the algorithm was repeated 10 times. The template with the highest average sliding window correlation time series across the 10 runs was chosen as the final result.

### Statistics and reproducibility

No calculations were used to predetermine sample size. It is important to note that this analysis was conducted at the within-subject level, with 1200 sampled time points per subject. Thus, the true ’sample size’ was substantially larger than 50 data points. Our analyses suggest that N=50 is sufficient to replicate the three spatiotemporal patterns across independent samples. No experimental conditions were randomized. No data was excluded from analysis. The Investigators were not blinded to allocation during experiments and outcome assessment.

## Acknowledgements

This work was supported by grants from the Canadian Institute for Advanced Research, a Gabelli Senior Scholar Award from the University of Miami, and R01MH107549 from the National Institute of Mental Health (NIMH) (to LQU), and an NIMH award (R03MH121668), and a NARSAD Young Investigator Award to (to JSN). BTTY was supported by the Singapore National Research Foundation (NRF) Fellowship (Class of 2017), the NUS Yong Loo Lin School of Medicine (NUHSRO/2020/124/TMR/LOA), the Singapore National Medical Research Council (NMRC) LCG (OFLCG19May-0035) and NMRC STaR (STaR20nov-0003).

## Author Contributions Statement

T.B. performed all analyses. S.K developed the original quasiperiodic pattern (QPP) algorithm used in this study. L.U, S.K, B.T.Y, D.B, J.N. and C.C assisted in the interpretation of analyses, conceptualization of the project, and writing of the manuscript. J.S assisted in the development and testing of the publicly-available Github repository that documents and stores analysis code.

## Data Availability

Data from the Human Connectome Project (HCP) is publicly available at http://www.humanconnectomeproject.org/data/. Instructions for accessing HCP data can be found at https://www.humanconnectome.org/. All metadata is provided at https://github.com/tsb46/BOLD_WAVES.

## Code Availability

All code for preprocessing and analysis is provided at https://github.com/tsb46/BOLD_WAVES.

